# FGF13 regulates cardiomyocyte impulse propagation via Cx43 trafficking independent of voltage-gated sodium channels

**DOI:** 10.1101/2025.07.23.666473

**Authors:** Lala Tanmoy Das, Mattia Malvezzi, Aravind Gade, Maiko Matsui, Margaret McKay, Eric Q. Wei, Matea J. Zelich, Keon Mazdisnian, Jared Kushner, Bi-xing Chen, Isabella DiStefano, Daniel Roybal, Lin Yang, Lisa Stoll, James C. Lo, Marian Kalocsay, Fadi G. Akar, Steven O. Marx, Geoffrey S. Pitt

## Abstract

**Background:** Fibroblast growth factor homologous factor (FHF) variants associate with arrhythmias. Although FHFs are best characterized as regulators of voltage gated sodium channel (VGSC) gating, recent studies suggest broader, non-VGSC-related functions, including regulation of Cx43 gap junctions and/or hemichannels, mechanisms that have generally been understudied or disregarded.

**Methods:** We assessed cardiac conduction and cardiomyocyte action potentials in mice with constitutive cardiac-specific *Fgf13* ablation (c*Fgf13^KO^*) while targeting Cx43 gap junctions and hemichannels pharmacologically. Using immunostaining and biochemistry, we characterized FGF13 regulation of Cx43 abundance and subcellular distribution. With proximity labeling proteomics, we investigated novel candidate mechanisms underlying FGF13 regulation of Cx43.

**Results:** FGF13 ablation prolonged the QRS and QT intervals on the surface electrocardiogram. Carbenoxolone, a Cx43 gap junction uncoupler, markedly prolonged the QRS duration leading to conduction system block in c*Fgf13^KO^* but not in WT mice. Optical mapping revealed markedly decreased conduction velocity (CV) during ventricular pacing. Microscopy revealed markedly perturbed trafficking of Cx43, reduced localization in the intercalated disc, and suggested decreased membrane Cx43 but increased Cx43 hemichannels in cardiomyocytes from c*Fgf13^KO^*mice. Resting membrane potential (RMP) was depolarized and APD50 was prolonged in c*Fgf13^KO^*cardiomyocytes. Both were restored towards wildtype (WT) values with Gap19 (a Cx43 hemichannel inhibitor), expression of FGF13, or expression of a mutant FGF13 incapable of binding to VGSCs, emphasizing VGSC-independent regulation by FGF13. To assess the functional impact of RMP depolarization, hearts were subjected to hypokalemia, which had no effect in WT hearts but fully rescued CV in c*Fgf13^KO^* hearts. Proteomic analyses revealed candidate roles for FGF13 in the regulation of vesicular-mediated transport. Biochemistry and immunocytochemistry showed that FGF13 ablation destabilized microtubules and reduced the expression of tubulins and MAP4, the major cardiac microtubule regulator.

**Conclusions:** FGF13 regulates microtubule-dependent trafficking and targeting of Cx43, thereby impacting cardiac impulse propagation via VGSC-independent mechanisms.

## INTRODUCTION

The fibroblast growth factor homologous factors (FHFs) are a subset of the fibroblast growth factors (FGFs) that are functionally distinct from canonical FGFs. Lacking signal sequences, FHFs (FGF11-FGF14) remain intracellular, are not secreted, do not bind to FGF receptors, nor function as growth factors ^1^. In the heart, FHFs have been best characterized as cytoplasmic interactors and regulators of voltage-gated sodium channels (VGSCs) and are linked to arrhythmogenesis ^2–8^. Human cardiomyocytes express both FGF12 and FGF13, with a FGF12 predominance ^2^. A variant in *FGF12* that perturbs the interaction with the main cardiac VGSC Na_V_1.5 has been associated with Brugada syndrome ^6^ and, conversely, a mutation in Na_V_1.5 that reduces the channel’s binding affinity for FGF12 has been associated with another inherited arrhythmia syndrome ^8^. More recently, a genome-wide association study for atrial fibrillation identified *FGF13* as a risk locus ^9^ and decreased left atrial expression of *FGF13* was associated with postoperative atrial fibrillation ^10^. Whether these FHF-associated arrhythmias arise solely from voltage-gated sodium channel dysfunction, however, is unclear. Attempts to define the full complement of effects mediated by FHFs and interpret the consequences of FHF variants on cardiac physiology and disease have been limited.

Connexin 43 (Cx43), the main connexin in ventricular cardiomyocytes, forms gap junctions at the intercalated discs (IDs), where VGSCs are concentrated, to allow for cell-cell electrical communications ^11^. In mice in which FGF13, the predominant cardiac FHF in rodents, was ablated globally, cardiac conduction shows increased sensitivity to gap junction blockers or reduced levels of Cx43 ^12^. While those effects were initially attributed solely to reduced sodium channel conductance resulting from *Fgf13* knockout, unmasked by pharmacological inhibition or genetic downregulation of Cx43 as a secondary hit, emerging evidence suggests that FGF13 also participate in additional cellular processes beyond VGSC regulation. ^13–16^ These observations prompted us to hypothesize in the current study that FGF13 *directly* modulates cardiac conduction through Cx43-dependent mechanisms. Building upon data showing that FGF13 stabilizes microtubules in the brain ^13^, we investigated whether FGF13 regulates intracellular trafficking of Cx43. Here, we identify a new role for FGF13 as a critical regulator of Cx43 trafficking in ventricular cardiomyocytes with consequences on cardiac conduction and show that this effect is independent of VGSC regulation.

## RESULTS

### Electrocardiographic analysis and optical mapping of cardiac-specific *Fgf13* knockout mice suggests roles beyond voltage-gated sodium channel regulation

To explore non-VGSC roles of the X-linked FGF13 in the heart, we generated constitutive, cardiac-specific *Fgf13* knockout (c*Fgf13^KO^*) mice by crossing females from a validated *Fgf13^fl/fl^* mouse line ^16^ with *Myh6-Cre^+^* male mice. Efficient knockout of FGF13 was confirmed by western blot and quantitative RT-PCR (**Supplemental Figure 1A-B**). Loss of FGF13 was not associated with a change in heart weight/body weight or gross morphology (**Supplemental Figure 1C-D**). Echocardiograms on conscious mice showed no differences in left ventricular function between the c*Fgf13^KO^* and wild-type (WT) control mice (**Supplemental Figure 1E-F**). However, ECGs showed that c*Fgf13^KO^*mice have longer QRS durations and QT intervals compared to controls (**Figure 1A-B**). These results suggest that, despite preserved gross cardiac anatomy and function, baseline electrical activation through the conduction system—and possibly the ventricular myocardium—is delayed by constitutive FGF13 knockout. This finding is consistent with prior observations in inducible *Fgf13* knockout ^16^ and after *Fgf13* knockdown ^2^, although not in a global *Fgf13* knockout model ^12^. The reason(s) for the different effects upon baseline cardiac conduction across models remain unclear, but because several studies showed that global *Fgf13* ablation is not viable^15,17–19^, we suspect that the reported global knockout mice retain residual *Fgf13* expression, potentially attenuating the observed conduction phenotype.

**Figure 1.**
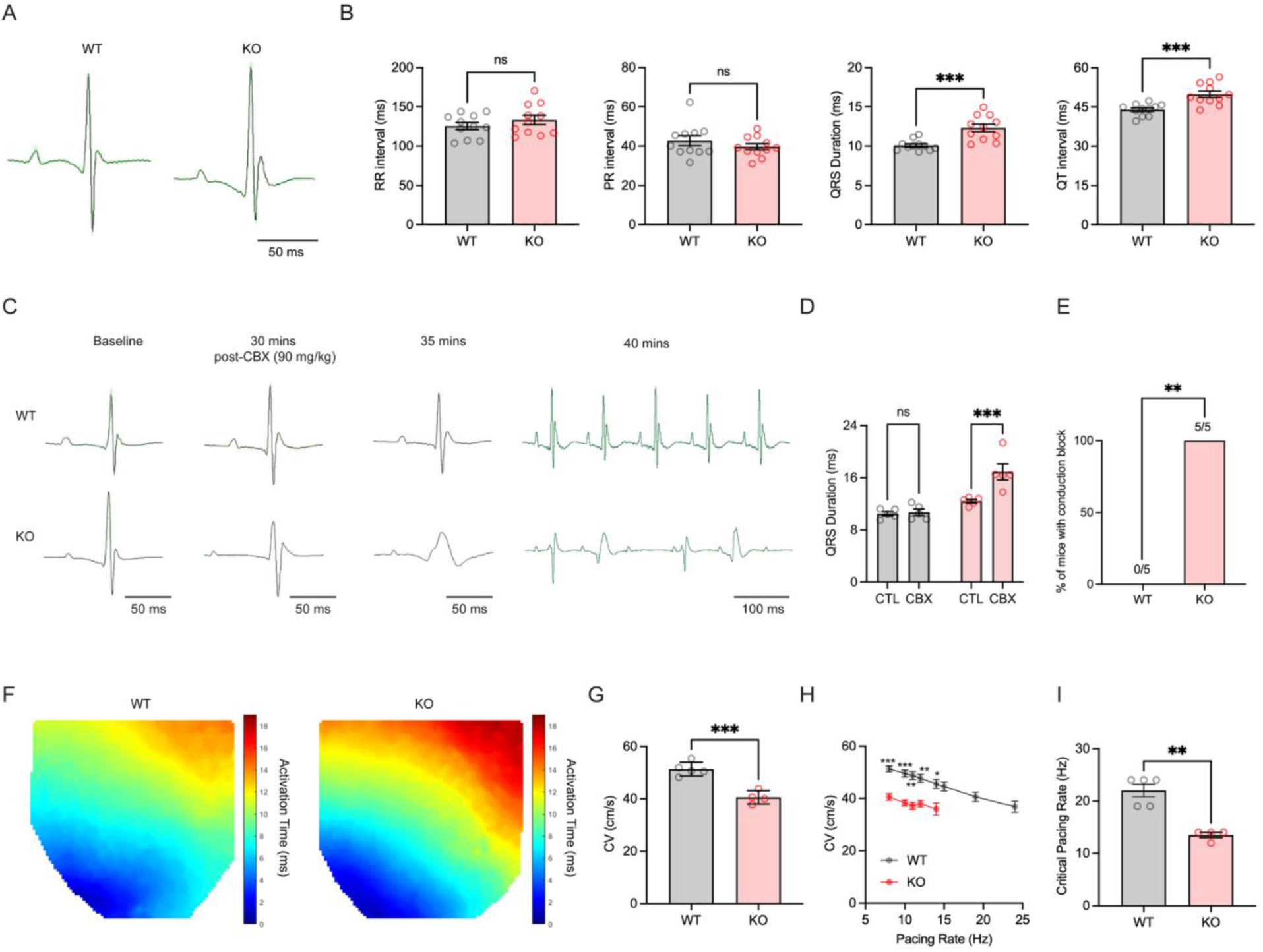
Cardiac-specific loss of FGF13 prolongs the QRS duration which is sensitive to gap junction blockade and slows conduction velocity. **A.** Representative lead I ECG traces from WT and c*Fgf13^KO^* mice at baseline. **B.** Summarized data (*n* = 11 mice per group) for RR interval (*p* = 0.31), PR interval (*p* = 0.31), QRS duration (*p* = 0.0008), and QT interval (*p* = 0.0006), 2-tailed, unpaired *t*-tests with Welch’s correction. **C.** Representative ECG traces of WT and KO mice at baseline, and 30, 35, 40 minutes after intraperitoneal injection of 90 mg/kg carbenoxolone [CBX] (*N* = 5 mice per group). **D.** Summarized data (*N* = 5 per group) for QRS duration at 30 minutes post-CBX injection, *p* = 0.0003 KO vs KO + CBX, 2-way ANOVA corrected for multiple comparisons. **E.** Quantification of conduction block occurrence after CBX administration. All KO mice (5/5) developed conduction block with 90 mg/kg carbenoxolone whereas WT littermates were unaffected, *p =* 0.0079, Fisher’s exact test. **F.** Representative isochrones demonstrating slower baseline conduction velocity (CV) in c*Fgf13^KO^* mouse hearts. **G.** CV (*N=*5 WT and 4 c*Fgf13^KO^* mice) at a pacing rate of 8 Hz, *p*=0.0005, unpaired *t-*test with Welch’s correction. **H.** Slowed CV in the c*Fgf13^KO^*hearts is significant at all pacing rates tested until capture in the c*Fgf13^KO^*group was lost. **I.** Critical pacing rate (the highest rate at which hearts maintained 1:1 capture), *p* = 0.0011, unpaired *t*-test with Welch’s correction. \*\**p* < 0.01, \*\*\**p* < 0.001, \*\*\*\**p* < 0.0001.

While the QRS prolongation in c*Fgf13^KO^* mice could be attributed to known effects on VGSC functions and/or availability attributed to FGF13 ^2,3,20^, previous data showing that FGF13 affects voltage-gated potassium channels ^16^ and contributes to cellular functions beyond regulation of VGSCs ^13–16^ prompted us to consider whether QRS prolongation reflects broader effects of FGF13 on ionic currents or cell to cell communication. Specifically, since the *Fgf13* global knockout mice showed increased sensitivity to pharmacological inhibition of Cx43 by the gap junction blocker carbenoxolone or its genetic knockdown in cardiomyocytes ^12^, we focused on Cx43 as a primary target of FGF13 regulation. We hypothesized that FGF13 altered Cx43 function independently from VGSCs. To test this, we first recorded surface ECGs in our c*Fgf13^KO^*mice and their WT controls after treatment with 90 mg/kg carbenoxolone, a dose at which ECG changes were detected in the whole-body knockout model. WT littermates were unaffected by carbenoxolone, but all c*Fgf13^KO^* mice showed not only progressive QRS prolongation but also high-degree atrioventricular block (**Figure 1C-E, Supplemental Figure 2**).

While QRS duration primarily reflects activation through the specialized conduction system, myocardial conduction can be directly measured using optical mapping during ex vivo ventricular pacing. Using this approach, we found that conduction velocity (CV) was significantly slower in c*Fgf13^KO^*ventricles compared to WT (**Figure 1F–G**). To further explore the basis for slowed CV, we assessed rate-dependent changes in conduction by varying the pacing frequency. At every pacing rate tested, including the slowest (8 Hz), CV remained consistently lower in c*Fgf13^KO^*hearts than in WT hearts (**Figure 1G–H**). Of note, the extent of CV slowing was comparable at both basal and rapid pacing rates, arguing against a role for impaired rate-dependent recovery from Na⁺ channel inactivation—one of the known consequences of FGF13 loss. Moreover, the critical pacing rate, the highest frequency at which 1:1 capture was maintained—was significantly reduced in cFgf13KO hearts (**Figure 1I**), indicating impaired conduction reserve.

Because the previously reported whole-body knockout model showed no QRS widening nor atrioventricular block even at a higher dose of carbenoxolone (120 mg/kg), those data suggest that the whole-body knockout mice were less sensitive to carbenoxolone than the c*Fgf13^KO^* mice studied here. We hypothesized that those whole-body knockout mice have a larger Cx43 reserve. Indeed, conduction block in those mice was only detected when those mice were also haploinsufficient for Cx43. The etiology of the larger Cx43 reserve in that model is not known but is consistent with our hypothesis that those mice have residual FGF13 and consequent reduced effects on Cx43. Our results thus raise the possibility that impaired Cx43-mediated conduction through gap junctions and/or Cx43 hemichannels contributes to the QRS widening, atrioventricular block, and slowed conduction velocity when *Fgf13* is eliminated.

### Constitutive *Fgf13* knockout hearts show altered Cx43 protein expression and spatial patterning

We next examined whether FGF13 ablation impacts Cx43 binding to Na_V_1.5. As they are both known direct Na_V_1.5 interactors ^5,21^, we reasoned that if the effect of FGF13 ablation on cardiac activation is due to perturbed VGSC modulation, then the Cx43-Na_V_1.5 interaction would remain unaltered. Instead, in an analysis of a previous dataset cataloging proteins co-immunoprecipitated with Na_V_1.5 in WT vs. c*Fgf13^KO^*hearts ^22^, we observed a 63% reduction (*P*=0.009) in Cx43 co-immunoprecipitated in c*Fgf13^KO^* hearts. This reduction in the amount of Cx43 co-immunoprecipitated with Na_V_1.5 in c*Fgf13^KO^* hearts suggests that less Cx43 is in complex with Na_V_1.5 in c*Fgf13^KO^* hearts, perhaps suggesting altered subcellular Cx43 localization.

We therefore systematically investigated the consequences of *Fgf13* knockout on Cx43 protein in ventricular tissue and in isolated ventricular cardiomyocytes. As expected, Cx43 in WT hearts was highly enriched at IDs, marked by N-cadherin. In tissue sections from c*Fgf13^KO^* hearts, however, we detected Cx43 in large puncta that did not co-localize with N-cadherin as reflected by a markedly reduced Spearman’s correlation coefficient (**Figure 2A-B**). N-cadherin, in contrast, was unaffected by loss of FGF13 (**Supplemental Figure 3A-D**). Much of the Cx43 signal in c*Fgf13^KO^* sections localized to the cardiomyocyte lateral membranes (marked by wheat germ agglutinin [WGA]) rather than to the IDs (**Figure 2C**), indicating a likely increased abundance of Cx43 hemichannels—Cx43 subunits along the cardiomyocyte lateral membranes that are not docked to other Cx43 subunits from adjacent cells and thus do not form typical gap junction channels ^23^.

**Figure 2.**
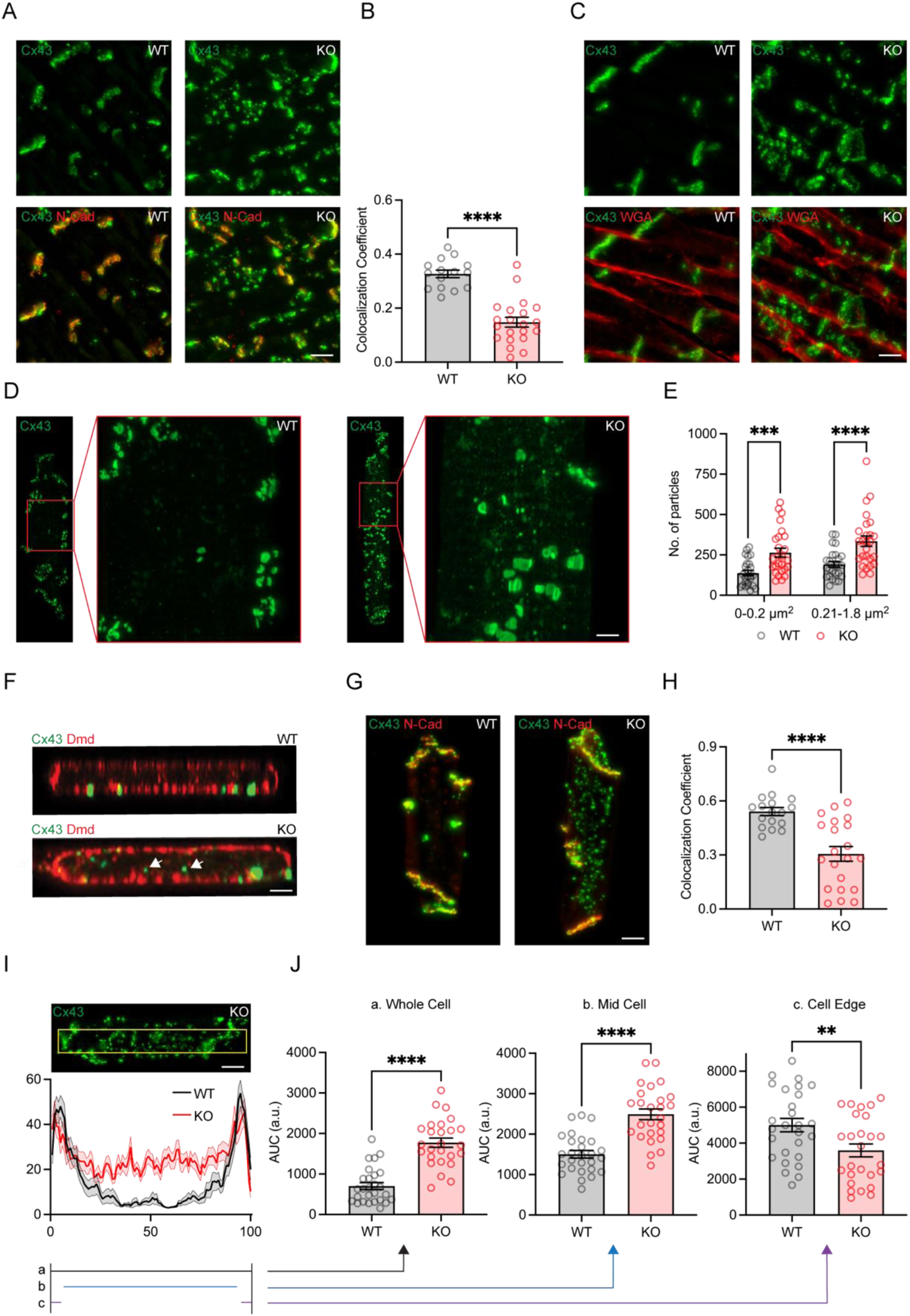
Loss of FGF13 perturbs Cx43 cellular distribution. **A.** Representative images from WT and c*Fgf13^KO^* paraffin-embedded ventricular tissue sections immunostained for Cx43 (green) and N-cadherin (red). Scale bar, 10 µm. **B.** Quantification of Spearman’s colocalization coefficient between Cx43 and N-cadherin from tissue sections in **A** (*N =* 3 mice per group, WT *n*= 15 sections, KO *n* = 21 sections), *p* < 0.0001, 2-tailed, unpaired *t*-test with Welch’s correction. **C.** Representative images from WT and c*Fgf13^KO^* paraffin-embedded ventricular tissue sections immunostained for Cx43 (green) and Wheat Germ Agglutinin [WGA] (red). Scale bar, 10 µm. **D.** Representative confocal images (z-stack max projection) of WT and c*Fgf13^KO^* isolated ventricular myocytes immunostained with Cx43 (green). Insets are magnified cellular areas showing different populations of Cx43 puncta. Scale bar, 10 µm. **E.** Quantification of the relative abundance of Cx43 secretory vesicles (0-0.2 mm^2^) and annular gap junctions (0.21-1.8 mm^2^) from confocal images of WT (gray) and KO (red) ventricular myocytes (*N* = 4 mice per group, WT *n* = 25 cells, KO *n* = 26 cells), *p* = 0.0004 for WT vs KO (0-0.2 mm^2^), *p* < 0.0001 for WT vs KO (0.21-1.8 mm^2^), 2-way ANOVA corrected for multiple comparisons. **F.** Representative orthogonal slices from confocal images of WT and KO isolated cardiomyocytes immunostained for Cx43 (green) and lateral membrane protein Dystrophin (Dmd, red). White arrows point to a non-lateral membrane bound Cx43 particle in a KO myocyte. Scale bar, 5 µm. **G.** Representative images of WT and c*Fgf13^KO^*isolated ventricular myocytes immunostained for Cx43 (green) and N-cadherin (red). Scale bar, 20 µm. **H.** Quantification of Spearman’s colocalization coefficient between Cx43 and N-cadherin from isolated myocytes (*N =* 3 mice per group, WT *n* = 17 cells, KO *n* = 21 cells), *p* < 0.0001, 2-tailed, unpaired *t*-test with Welch’s correction. **I.** Pixel intensity analysis across longitudinal axes of WT and c*Fgf13^KO^* cardiomyocytes (*N* = 4 mice per group, WT *n* = 28 cells, KO *n* = 26 cells). Cell distance and intensity are normalized to a WT cell. **J.** Cx43 pixel intensity along the cell is quantified using Area Under the Curve (AUC) in three groups: Whole-cell (black line), Mid cell (10 – 90 normalized distance units, blue line), Cell edge (0-5 and 95-100 normalized distance units, purple line). \**p* < 0.05, \*\**p* < 0.01, \*\*\**p* < 0.001, \*\*\*\**p* < 0.0001.

We extended our analyses to freshly isolated single ventricular cardiomyocytes, which offer a uniform orientation for accurate quantification. Confocal microscopy revealed two populations of Cx43 punctae (**Figure 2D-E**) consistent with previous observations ^24^: large punctae, such as those observed in tissue and that display characteristics and morphology consistent with annular gap junctions after endocytosis; and small punctae that likely represent Cx43-loaded vesicles trafficking along microtubules. Using those previously established metrics^24^ we quantified the relative abundance of likely annular gap junctions (defined as ≥0.5 µm diameter and average area of 1.6±0.3 µm^2^) and likely Cx43-loaded vesicles (defined as ≤150 nm in diameter) and found that both populations were larger in c*Fgf13^KO^* cardiomyocytes relative to controls. Moreover, many of the Cx43 punctae in c*Fgf13^KO^* cardiomyocytes were within the cytoplasm and not associated with the sarcolemma, while Cx43 punctae in WT control cells were generally localized at the cell membrane (**Figure 2F** and **Supplemental Figure 4**). We also observed many large Cx43 punctae scattered throughout c*Fgf13^KO^* isolated cells, while in WT isolated cardiomyocytes Cx43 punctae predominantly colocalized with N-cadherin at the IDs (**Figure 2G-H**). There were also more Cx43 punctae along the lateral membrane in c*Fgf13^KO^*cardiomyocytes (likely hemichannels) than in WT cells. While acute cardiomyocyte isolation could perturb the localization of ID components, the consistent genotype-specific patterns and the abnormal Cx43 distribution in c*Fgf13^KO^* cells were similar to what we observed in tissue, providing validation for the isolated cardiomyocyte results. We therefore exploited the acutely isolated cells to quantify the distribution of Cx43 along the long axis of cardiomyocytes. Compared to WT cells, Cx43 signal intensity in c*Fgf13^KO^* ventricular cardiomyocytes was larger between the cell ends (*i.e.*, between the IDs) (**Figure 2I-J**) and smaller at the cell ends.

Immunohistochemistry and immunocytochemistry for Cx43 in tissue and cells revealed not only a genotype difference in Cx43 spatial patterning but also suggested increased Cx43 abundance in c*Fgf13^KO^* myocytes. We therefore quantified the % area of the cardiomyocyte positive for Cx43 immunofluorescence (**Figure 3A-B**), which was greater in c*Fgf13^KO^* cardiomyocytes compared to WT cardiomyocytes. Consistent with this observation, we also quantified the total amount of Cx43 by western blot in lysates from isolated ventricular myocytes and found that Cx43 was 43 ± 9.4% (*P*<0.01) higher in c*Fgf13^KO^* cells (**Figure 3C-D**). Although Cx43 total protein was increased, surface biotinylation performed with isolated cardiomyocytes showed that Cx43 was reduced at the cell surface by 28 ± 6% (*P*<0.01) in c*Fgf13^KO^* cells (**Figure 3E-F**). This finding is consistent with our observation that Cx43 punctae in c*Fgf13^KO^* cardiomyocytes were generally within the cytoplasm and not at the cell surface (see **Figure 2F** and **Supplemental Figure 4**). Thus, we conclude that cardiac-specific, constitutive knockout of *Fgf13* increases Cx43 total protein yet decreases Cx43 abundance at the surface and perturbs Cx43 localization. To determine if the greater Cx43 protein abundance in c*Fgf13^KO^* cardiomyocytes resulted from increased Cx43 transcription, we performed qPCR. We did not detect a difference between WT and c*Fgf13^KO^* (**Supplemental Figure 5A**). We also analyzed Cx43 stability using cycloheximide (100 µg/ml) applied to isolated cardiomyocytes to block translation. After three hours, we observed no difference in the amount of Cx43 degradation between WT and c*Fgf13^KO^*cells (**Figure 3G-H**), suggesting alternative mechanisms.

**Figure 3.**
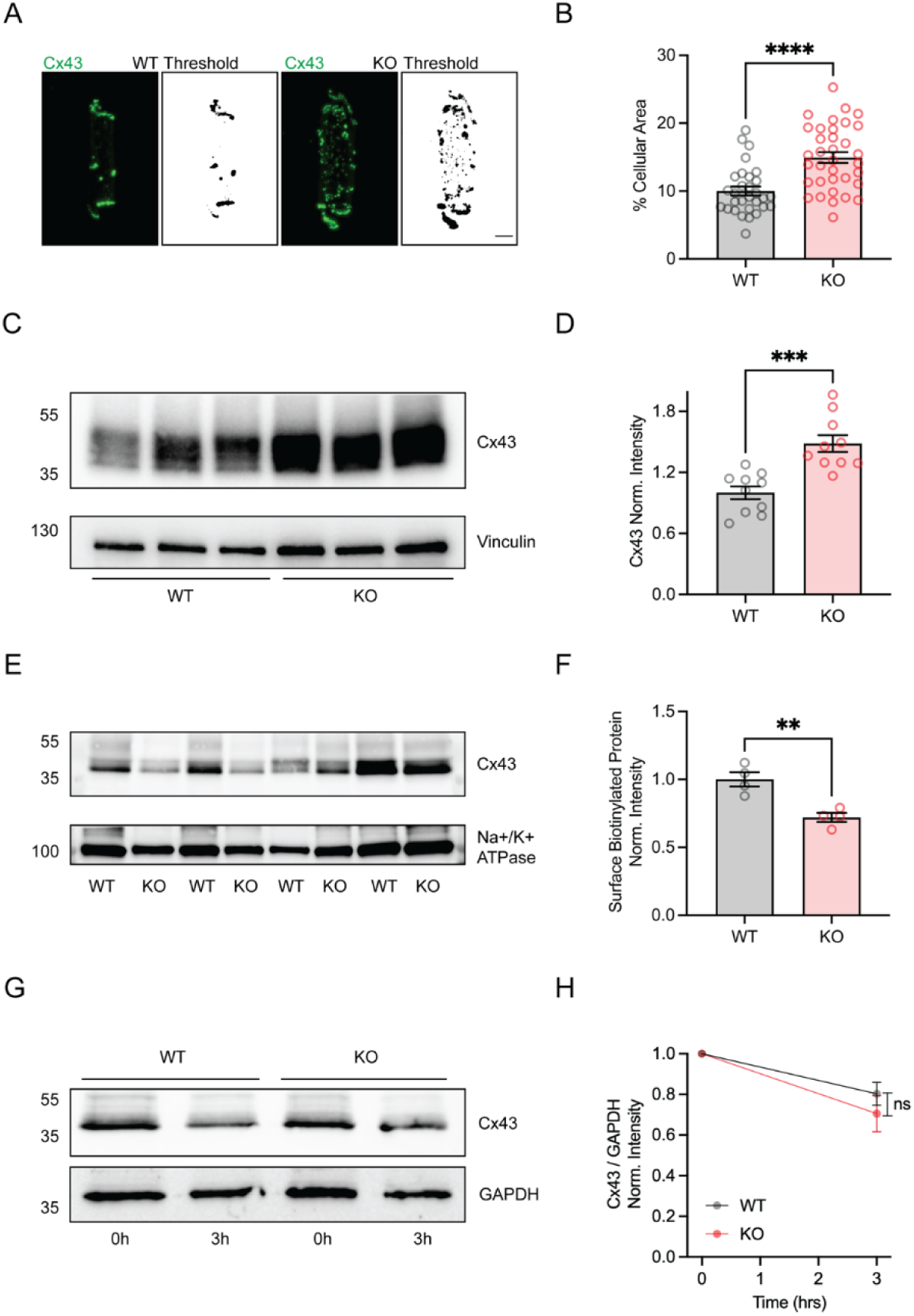
Loss of FGF13 alters Cx43 protein expression. **A.** Representative images of WT and c*Fgf13^KO^* isolated cardiomyocytes immunostained with Cx43 (green) and respective 8-bit images with thresholding. Scale bar, 10 µm. **B.** Quantification of Cx43 expression from thresholded images approximated as % Cellular Area (*N* = 4 mice per group, WT *n* = 30 cells, KO *n* = 35 cells), *p* < 0.0001, 2-tailed, unpaired *t*-test with Welch’s correction. **C.** Western blot (WB) of isolated ventricular cardiomyocyte lysates (*N* = 3 mice per group) immunoblotted for Cx43 and Vinculin. **D.** Western blot densitometric analysis of Cx43 protein expression normalized to Vinculin from isolated ventricular cardiomyocytes (*N* = 10 mice per group), 2-tailed, unpaired *t*-test with Welch’s correction, *p* = 0.0003. **E.** Western blot of Cx43 and Na+/K+-ATPase (membrane protein loading control) biotinylated fractions (*N* = 4 mice per group). **F.** Densitometric analysis of biotinylated Cx43 normalized to biotinylated Na+/K+-ATPase (*N =* 4 per group), 2-tailed, unpaired *t*-test with Welch’s correction, *p* = 0.006. **G.** Representative western blot of untreated isolated ventricular cardiomyocyte lysates (0h) and 3-hour cycloheximide-treated (100 ug/mL) isolated ventricular cardiomyocyte lysates (3h) immunoblotted for Cx43 and GAPDH. **H.** Densitometric analysis of untreated and 3h-cycloheximide treated cardiomyocyte lysates immunoblotted for Cx43 and GAPDH (*N* = 4 mice per group), *p* = 0.22, 2-way ANOVA corrected for multiple comparisons. \**p* < 0.05, \*\**p* < 0.01, \*\*\**p* < 0.001, \*\*\*\**p* < 0.0001.

### Analysis of action potentials in isolated cardiomyocytes suggests *Fgf13* knockout increases Cx43 hemichannels

Recent data show that disease states like heart failure lead to an increase in Cx43 hemichannels (upon pathological remodeling) and that the altered distribution and activity of hemichannels affect cardiomyocyte excitability and contribute to arrhythmias ^25^. Additional data from dystrophic mouse cardiomyocytes show that mislocalized and remodeled Cx43 hemichannels contribute to a depolarized resting membrane potential ^25–27^ that can be hyperpolarized upon addition of the Cx43 hemichannel blocker, Gap19 ^26,27^. Furthermore, pathological remodeling of Cx43 hemichannels can prolong the action potential duration, which can be reduced to nearly WT levels with the addition of Gap19 ^27^. Overall, these studies implicate a critical role of Cx43 hemichannels in electrophysiological properties of cardiomyocytes.

Because we observed significant lateralization of Cx43 in tissue and cardiomyocytes from c*Fgf13^KO^* hearts, we tested for the contributions of increased Cx43 hemichannels in c*Fgf13^KO^*mice by recording action potentials (APs) in isolated ventricular cardiomyocytes with the perforated patch technique. Consistent with previous data from an inducible knockout model, AP amplitude in c*Fgf13^KO^*myocytes was significantly reduced compared to controls, while rise time and APD50 (**Figure 4C-D**) were longer in c*Fgf13^KO^* myocytes compared to WT, which can be attributed at least in part to a hyperpolarizing shift in the steady state inactivation V_1/2_ of VGSC currents as observed in previous reports ^2,16,28–30^.

**Figure 4.**
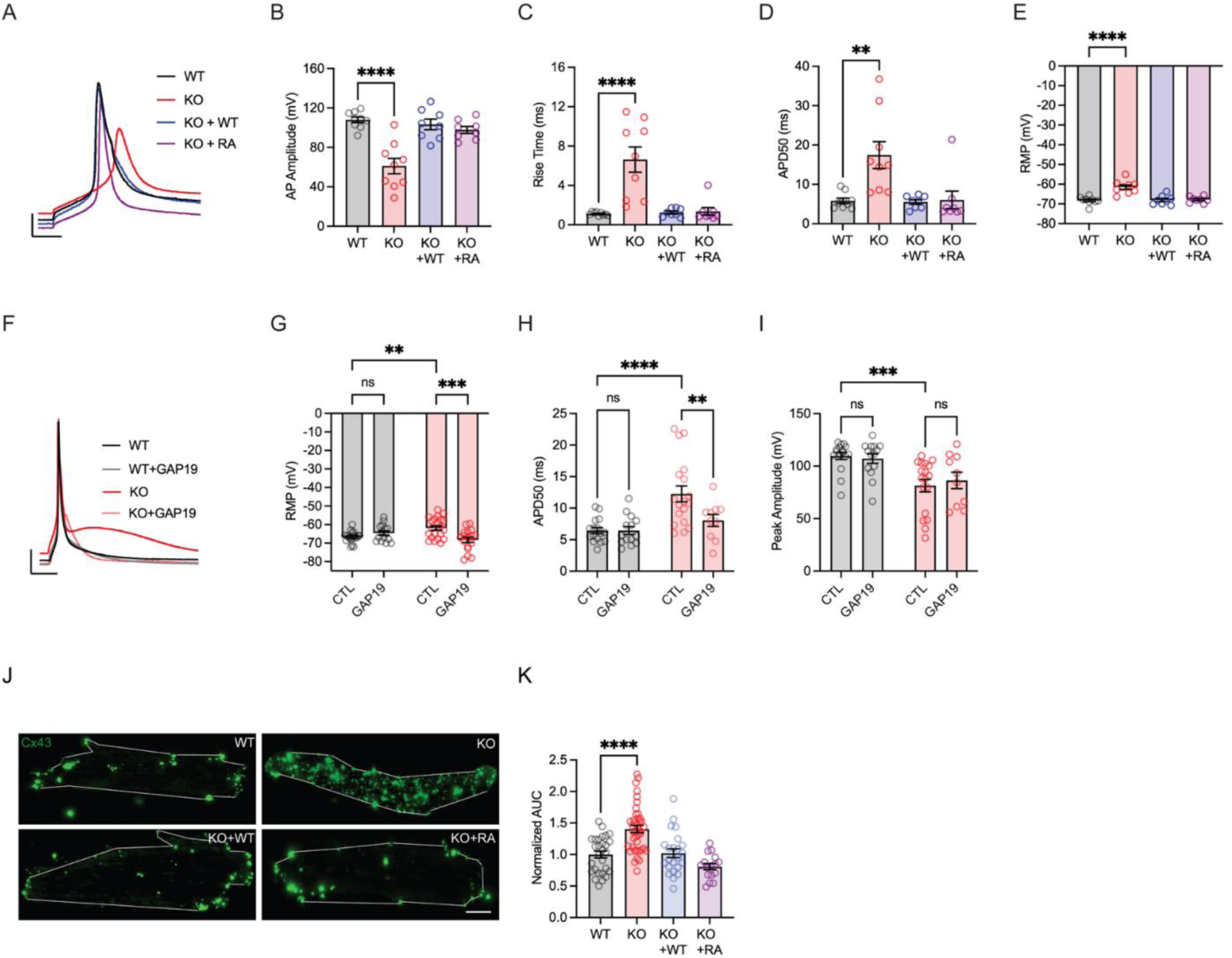
*Fgf13* knockout increases aberrant Cx43 hemichannel contributions to action potentials. **A**. Representative AP traces of 24-48 hour cultured ventricular cardiomyocytes using perforated patch clamp protocol from the following groups: WT (black), c*Fgf13^KO^* (red), c*Fgf13^KO^* + WT FGF13 AAV-mediated re-expression (blue), and c*Fgf13^KO^*+ FGF13^R/A^ AAV-mediated re-expression (purple). Scale bar: x-axis 10 ms, y-axis 20 mV. **B.** Summarized data for AP Amplitude (WT *N =* 3 mice, *n =* 9 cells, KO *N* = 3 mice, *n* = 9 cells, KO + WT *N* = 3 mice, *n* = 8 cells, KO + FGF13^R/A^ *N* = 3 mice, *n* = 8 cells; *p* < 0.0001 for WT vs KO only). **C**. Summarized data for AP rise time (WT *N* = 3 mice, *n =* 9 cells, KO *N* = 3 mice, *n* = 9 cells, KO + WT FGF13 *N* = 3 mice, *n* = 8 cells, KO + FGF13^R/A^ *N* = 3 mice, *n* = 8 cells; *p* < 0.0001 for WT vs KO only). **D.** Summarized data for APD50 (WT *N* = 3 mice, *n =* 9 cells, KO *N* = 3 mice, *n* = 9 cells, KO + WT FGF13 *N* = 3 mice, *n* = 8 cells, KO + FGF13^R/A^ *N* = 3 mice, *n* = 8 cells; *p* < 0.0001 for WT vs KO only). **E.** Summarized data for resting membrane potential (RMP, WT *N* = 3 mice, *n* = 9 cells, KO *N* = 3 mice, *n* = 9 cells, KO + WT FGF13 *N* = 3 mice, *n* = 8 cells, KO + FGF13^R/A^ *N* = 3 mice, *n* = 8 cells, *p* < 0.0001 for WT vs KO only, one-way ANOVA corrected for multiple comparisons. **F.** Representative action potential (AP) traces (whole-cell patch clamp) of isolated myocytes injected with a 200-pA current pulse for 50 milliseconds. WT (black), WT + GAP19 (gray), KO (red), and KO + GAP19 (pink). Scale bar: x-axis 50 ms, y-axis 20 mV **G.** Summarized data (*N* = 3 mice per group) for RMP in control (CTL) and GAP19-treated WT and c*Fgf13^KO^* isolated cardiomyocytes (WT *n =* 19 cells, WT + GAP19 *n* = 15 cells, KO *n* = 21 cells, KO + GAP19 *n* = 17 cells), 2-way ANOVA corrected for multiple comparisons, *p* = 0.0044 KO vs KO + GAP19 cells. **H.** Summarized data (*N* = 3 mice per group) for action potential duration at 50% (APD50, WT *n =* 16 cells, WT + GAP19 *n* = 14 cells, KO *n* = 18 cells, KO + GAP19 *n* = 10 cells), 2-way ANOVA corrected for multiple comparisons, *p* < 0.0001 WT vs KO cells, *p* = 0.004 KO vs KO + GAP19 cells. **I.** Summarized data (*n* = 4 mice per group) of Peak Amplitude (WT *n =* 16 cells, WT + GAP19 *n* = 14 cells, KO *n* = 18 cells, KO + GAP19 *n* = 10 cells, 2-way ANOVA corrected for multiple comparisons, *p* = 0.0002 for WT vs KO). **J.** Representative images of WT, c*Fgf13^KO^*, c*Fgf13^KO^* + WT FGF13, c*Fgf13^KO^* + FGF13^R/A^ ventricular myocytes cultured for 24-48 hours and immunostained with Cx43 (green). Scale bar, 10 µm. **K.** Cx43 pixel intensity analysis and normalized AUC (*n =* 3 mice per group, WT *n* = 28 cells, KO *n* = 41 cells, KO + WT FGF13 *n* = 23 cells, KO + FGF13^R/A^ *n* = 16 cells), 2-way ANOVA corrected for multiple comparisons, *p* = 0.0001 for WT vs KO only).

We also observed a more depolarized resting membrane potential (RMP) in cardiomyocytes isolated from c*Fgf13^KO^* cardiomyocytes than in WT cardiomyocytes, a parameter unlikely to be affected by VGSCs. To test that hypothesis definitively, we exploited a strategy to dissociate the contribution of FGF13 regulation of VGSCs by replacing WT FGF13 with a mutant version (Arg120Ala, “FGF13^R/A^”) unable to bind to Na_V_1.5 ^22,31^. We infected cardiomyocytes isolated from c*Fgf13^KO^* mice with adeno-associated viruses (AAV) expressing either WT FGF13, FGF13^R/A^, or GFP as a control. After 24-48 hours in culture to allow expression from the virus (all groups above were cultured in parallel to allow direct comparisons), we recorded RMPs and observed that addition of either FGF13 or FGF13^R/A^ hyperpolarized RMP to values similar to WT (**Figure 4E**).

We therefore hypothesized that the depolarized RMP in c*Fgf13^KO^* cardiomyocytes derived at least in part from increased Cx43 hemichannels, which we tested by adding the Cx43 hemichannel blocker, Gap19. Treatment with Gap19 requires intracellular application, so we utilized the whole-cell patch clamp technique to dialyze Gap19 via the patch pipette. The addition of Gap19 to the internal pipette solution hyperpolarized the resting membrane potential of c*Fgf13^KO^*cells but did not affect WT cardiomyocytes (**Figure 4F-G**), supporting our hypothesis that excess inward depolarizing current through Cx43 hemichannels (in the absence of Gap19) depolarized the resting membrane potential in c*Fgf13^KO^*cardiomyocytes.

Having established that excess Cx43 hemichannels contributed to the depolarized RMP in c*Fgf13^KO^* cardiomyocytes, we asked if these hemichannels affected other AP characteristics. We calculated the APD50, which showed an increased duration in c*Fgf13^KO^* cells compared to WT cells (**Figure 4H**). These results provide additional support for our hypothesis that excess hemichannels in c*Fgf13^KO^* cardiomyocytes affect AP properties and are consistent with a recent study showing that Cx43 hemichannels contribute to AP repolarization in isolated cardiomyocytes, especially in disease states in which hemichannels are more abundant ^25^. Further, we examined the AP peak amplitude, a measure that depends mainly on Na_V_1.5 channels (rather than Cx43 gap junctions). While in cells isolated from WT mice the peak amplitude was higher than in cells from *Fgf13^KO^* mice, addition of Gap19 to either WT or c*Fgf13^KO^* cells had no effect on the AP amplitude (**Figure 4I**), suggesting that Gap19 does not affect parameters regulated mainly by VGSCs. Moreover, we observed that expression of either FGF13 or FGF13^R/A^ in c*Fgf13^KO^* cardiomyocytes ameliorated the observed abnormal Cx43 distribution, leaving Cx43 mainly localized to cell edges while markedly reducing scattered punctae throughout the cytoplasm (**Figure 4J-K**).

We next hypothesized that the depolarized RMP resulting from increased Cx43 hemichannel activity in c*Fgf13^KO^* hearts impairs CV, likely by reducing Na_V_1.5 channel availability. To test this, we repeated CV measurements under low extracellular potassium (2 mM K⁺) conditions, which hyperpolarize the RMP by approximately 20 mV. If conduction slowing under physiological K⁺ (4 mM) in the c*Fgf13^KO^* hearts is due to RMP-mediated VGSC inactivation, then hyperpolarization should restore VGSC availability and normalize CV.

Indeed, under low K⁺ conditions, CV in c*Fgf13^KO^* hearts significantly increased across all pacing rates and became indistinguishable from WT hearts (**Figure 5A–B**). Similarly, the critical pacing cycle rate was normalized in c*Fgf13^KO^* hearts and matched values observed in WT controls (**Figure 5C**). These findings strongly support the conclusion that RMP depolarization, driven by excess Cx43 hemichannel activity, reduces VGSC availability and contributes to slowed conduction in c*Fgf13^KO^*hearts. The restoration of CV under hyperpolarized conditions reinforces the functional link between RMP and myocardial conduction defects in this model and contrasts sharply with models in which slowed conduction arises from impaired recovery of Na⁺ channels from inactivation, conditions under which hypokalemia typically worsens, rather than improves, conduction, especially during rapid pacing.

**Figure 5.**
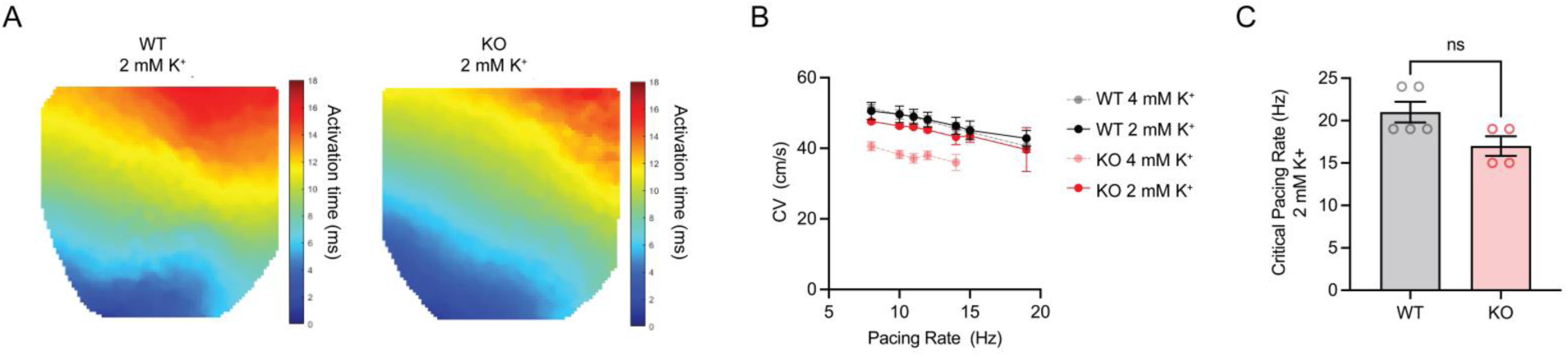
Hyperpolarization of resting membrane potential by low potassium restores conduction velocity in c*Fgf13^KO^* hearts. **A**. Representative isochrones showing baseline conduction velocity (CV) in WT and c*Fgf13^KO^* mouse hearts with 2 mM extracellular K^+^. **B.** CV in the c*Fgf13^KO^* hearts is accelerated in 2 mM K^+^, not different from CV in WT hearts at all pacing rates tested. Data from normal K^+^ (4 mM) repeated from Figure 1 for comparison. **C.** Critical pacing rate is not different between WT and c*Fgf13^KO^*with 2 mM extracellular K^+^.

That the Na_V_1.5 binding incompetent FGF13^R/A^ restored Cx43 co-localization further suggested that at least some FGF13 functions are independent of its interactions with Na_V_1.5. To test our hypothesis, we queried the relative Na_V_1.5-free fraction of FGF13 in cardiomyocytes. Immunoprecipitation of Na_V_1.5 successfully depleted 80% of the channel from cardiomyocyte lysates but only about 35% of FGF13 (**Figure 6A-B**), suggesting a stoichiometric excess of FGF13 compared to Na_V_1.5. Thus, a significant fraction of cardiomyocyte FGF13 does not interact with Na_V_1.5 and likely has other roles in cardiomyocytes, as implied by these results.

**Figure 6.**
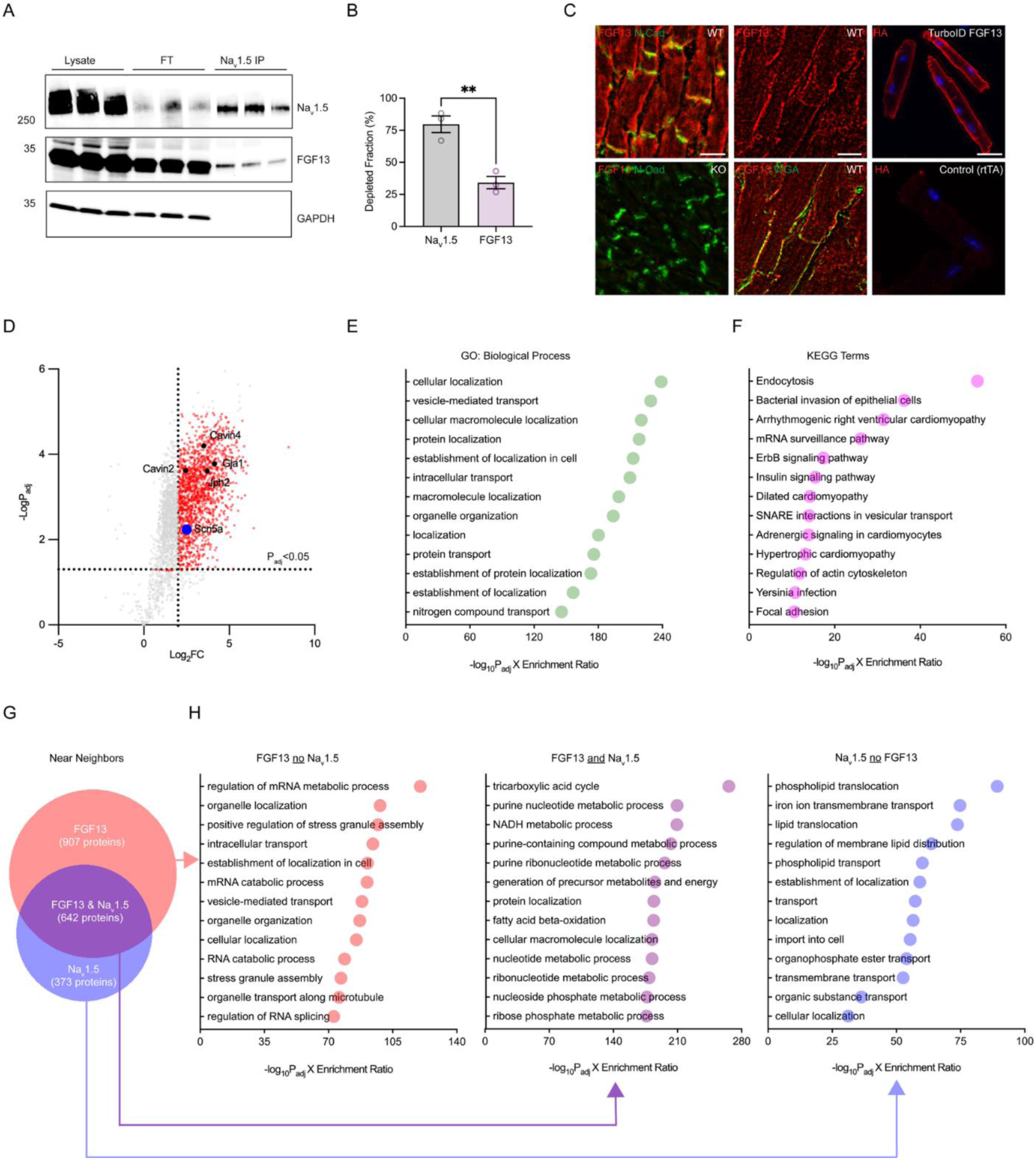
FGF13 protein neighbors are associated with non-VGSC binding functions such as vesicle-mediated transport. **A.** Immunoblot of WT heart lysates (*n* = 3 mice), SCN5A coimmunoprecipitated fractions, and corresponding flowthrough (FT) fractions probed for Na_v_1.5, FGF13, and GAPDH. **B.** Quantification of Nav1.5 CoIP shows that only about 35% of the total FGF13 is in complex. FT = Flowthrough. **C.** Left and middle panels are magnified images of left ventricular cryosections immunostained for FGF13 (red), N-cadherin (green) and wheat germ agglutinin [WGA] (green). Scale bar, 10 µm. Right panel shows isolated ventricular myocytes with FGF13 TurboID and control myocytes immunostained for hemagglutinin (HA). Scale bar, 20 µm. **D.** Volcano plot of significantly enriched terms from FGF13 TurboID proximity proteomics (P_adj_ < 0.05, Log_2_FC > 1). SCN5A, a known FGF13 interactor, is highlighted in a blue circle. Other proteins previously shown to coimmunoprecipitate with FGF13 are also indicated including Cavin4, Cavin2, and Junctophilin-2 (Jph2). Cx43 (Gja1) is also a significantly enriched FGF13 neighbor. **E.** Top 13 Gene Ontology (GO) terms for Biological Process from FGF13 TurboID gene set enrichment analysis (Log_2_FC > 2.0, P_adj_ < 0.05). x axis shows the product of -log_10_P_adj_ and Enrichment Ratio. **F.** Top 13 KEGG terms from FGF13 TurboID gene set enrichment analysis (Log_2_FC > 2.0, P_adj_ < 0.05). *x axis* shows the product of -log_10_P_adj_ and Enrichment Ratio. **G.** Venn diagram of FGF13 TurboID terms (Log_2_FC > 2.0, P_adj_ < 0.05) and SCN5A TurboID terms (Log_2_FC > 2.0, P < 0.05). **H.** Top 13 Biological Process GO terms from **G** separated into 3 categories: FGF13 only no Na_v_1.5 (salmon), Na_v_1.5 only no FGF13 (purple), FGF13 and Na_v_1.5 intersection (blue), *x axis* shows the product of -log_10_P_adj_ and Enrichment Ratio.

### FGF13 protein neighbors suggest a role in vesicle-mediated transport of Cx43

To investigate how FGF13 affects Cx43 localization within cardiomyocytes and targeting to the IDs, we performed an unbiased screen for candidate FGF13 near neighbors. We used protein proximity labeling in whole hearts from transgenic mice expressing FGF13 fused to the biotin ligase TurboID ^32^ and a hemagglutinin (HA) tag. We first validated that the subcellular localization of transgenic TurboID-FGF13 mimics endogenous FGF13 by visualizing the TurboID-FGF13 via HA immunofluorescence in isolated ventricular myocytes. Similar to endogenous FGF13 immunostaining in control tissue sections (**Figure 6C**), HA immunofluorescence showed enhanced signal along the lateral membrane and at the IDs (marked by N-cadherin). Biotinylation efficiency of the TurboID-FGF13 in mice injected with biotin was confirmed by western blot with streptavidin-HRP, which showed intense signals over a large range of molecular weights compared to control mice injected with 10% DMSO (n=3 mice, each), for which only endogenously biotinylated carboxylases were detected (**Supplemental Figure 6A**). Biotin-labeled proteins were captured by streptavidin and analyzed by semi-quantitative mass spectrometry. Compared to control samples without biotin, we identified 1549 unique proteins enriched in the FGF13 neighborhood (Log_2_ fold change [Log_2_FC]>2.0 and -log_10_P_adj_>1.30 [P<0.05], **Supplemental Dataset 1**). As expected, we identified Na_V_1.5 (*Scn5a*, Log_2_FC=2.50, -log_10_P_adj_=2.08) (**Figure 6D**) and a set of proteins previously shown to coimmunoprecipitate with FGF13 from heart lysates ^15,33^ such as junctophilin-2 (*Jph2*, Log_2_FC=3.71, -log_10_P_adj_=2.88), cavin-2 (*Cavin2*, Log_2_FC=2.45, - log_10_P_adj_=2.88), and cavin-4 (*Cavin4*, Log_2_FC=3.50, -log_10_P_adj_=2.99). We also identified Cx43 (*Gja1*, Log_2_FC=4.13, -log_10_P_adj_=2.92), providing further support for our hypothesis that FGF13 regulates the trafficking and targeting of Cx43.

We performed gene set enrichment analysis on the entire set and ranked the gene ontology (GO) terms by the product of their Enrichment Ratio (Gene Ratio / Effective_domain_size; see **Methods**) and adjusted P-value, which revealed that among the GO biological processes (**Figure 6E**) the top 2 terms were “cellular localization” (GO:0051641) and “vesicle-mediated transport” (GO:0016192). We also performed a gene enrichment analysis using the Kyoto Encyclopedia of Genes and Genomes (KEGG) (**Figure 6F**) and found that the top term was “endocytosis” (KEGG:04144). Together, these analyses implicate FGF13’s role in bidirectional (anterograde and retrograde) transport processes.

Since an overarching goal was to identify FGF13 functions that are distinct from regulation of Na_V_1.5, we performed a separate protein proximity labeling experiment with TurboID fused to Na_v_1.5, allowing us to focus on the non-overlapping set of unique FGF13 near neighbors (**Supplemental Dataset 2**). The Na_V_1.5 TurboID experiment, performed using tandem mass tags (TMT), identified 1015 proteins (Log_2_FC>2.0, P<0.05), of which 642 were also contained in the FGF13 dataset, in which label-free quantification was applied. Among the FGF13 near neighbors, 907 proteins (58.5%) were absent in the Na_v_1.5 dataset (**Figure 6G-H**), on which we then performed gene set enrichment analysis. Among the top hits were “intracellular transport” (GO:0046907), “vesicle-mediated transport” (GO:0016192), and multiple terms associated with microtubules (**Supplemental Figure 7A**). Gene set enrichment analysis of the 373 proteins unique to the Na_V_1.5 dataset did not return any of these or related terms (**Figure 6G-H**). Thus, we focused additional studies on querying whether FGF13 regulates Cx43 via mechanisms related to vesicular transport.

### FGF13-mediated microtubule instability alters Cx43 localization pattern

Building on these analyses from the TurboID data sets and on previous data suggesting that FGF13 can stabilize microtubules ^13^, including in cardiomyocytes ^34^, we queried whether FGF13 affected microtubules as a mechanism to regulate Cx43 by first exploring an interaction between FGF13 and microtubules. In lysates from WT cardiomyocytes, alpha tubulin (but not beta tubulin) co-immunoprecipitated with FGF13 (**Figure 7A**), consistent with a recent study ^34^. We then performed immunohistochemistry to evaluate alpha tubulin in tissue sections and found that the density and intensity of alpha tubulin differed between control and KO hearts (**Figure 7B, Supplemental Figure 7B**). Given the constraints in achieving consistent tissue orientation for quantitative analyses, we exploited acutely isolated ventricular cardiomyocytes. Using a thresholding protocol, we found that c*Fgf13^KO^*cells had lower tubulin density compared to WT cardiomyocytes (**Figure 7C-D**). Further, we observed an increased immunofluorescence intensity of acetylated alpha tubulin, a marker of microtubule stability ^35^, in cardiomyocytes isolated from WT compared to c*Fgf13^KO^* mice (**Figure 7E-F**). Infection of c*Fgf13^KO^* cardiomyocytes with an AAV expressing FGF13 increased the acetylated tubulin immunofluorescence intensity, suggesting that re-expression of FGF13 plays a dynamic role in tubulin acetylation and microtubule stability (**Figure 7E-F**). Infection of c*Fgf13^KO^* cardiomyocytes with the Na_V_1.5 binding-incompetent mutant FGF13^R/A^ also increased the acetylated tubulin immunofluorescence intensity, emphasizing that the effect on acetylated tubulin is independent of FGF13’s interaction with Na_V_1.5 (**Figure 7E-F**). We also quantified the relative fraction of acetylated alpha tubulin and total alpha tubulin by western blot. We observed a significant decrease in both in c*Fgf13^KO^* cardiomyocytes compared to WT cells (**Figure 7G-I**). Beta tubulin protein was similarly reduced in c*Fgf13^KO^*heart lysates compared to WT (**Figure 7J**). Immunofluorescence also showed decreased signal for beta tubulin in isolated c*Fgf13^KO^* cardiomyocytes (**Figure 7K-L**). Further, qRT-PCR revealed that transcripts for *Tuba4a*, one of the most highly expressed isotypes in the heart ^36^, were significantly decreased (**Supplemental Figure 5B**). Thus, FGF13 knockout not only decreases microtubule stability but also reduces tubulin protein, suggesting that FGF13 exerts multiple regulatory controls on tubulins and microtubules.

**Figure 7.**
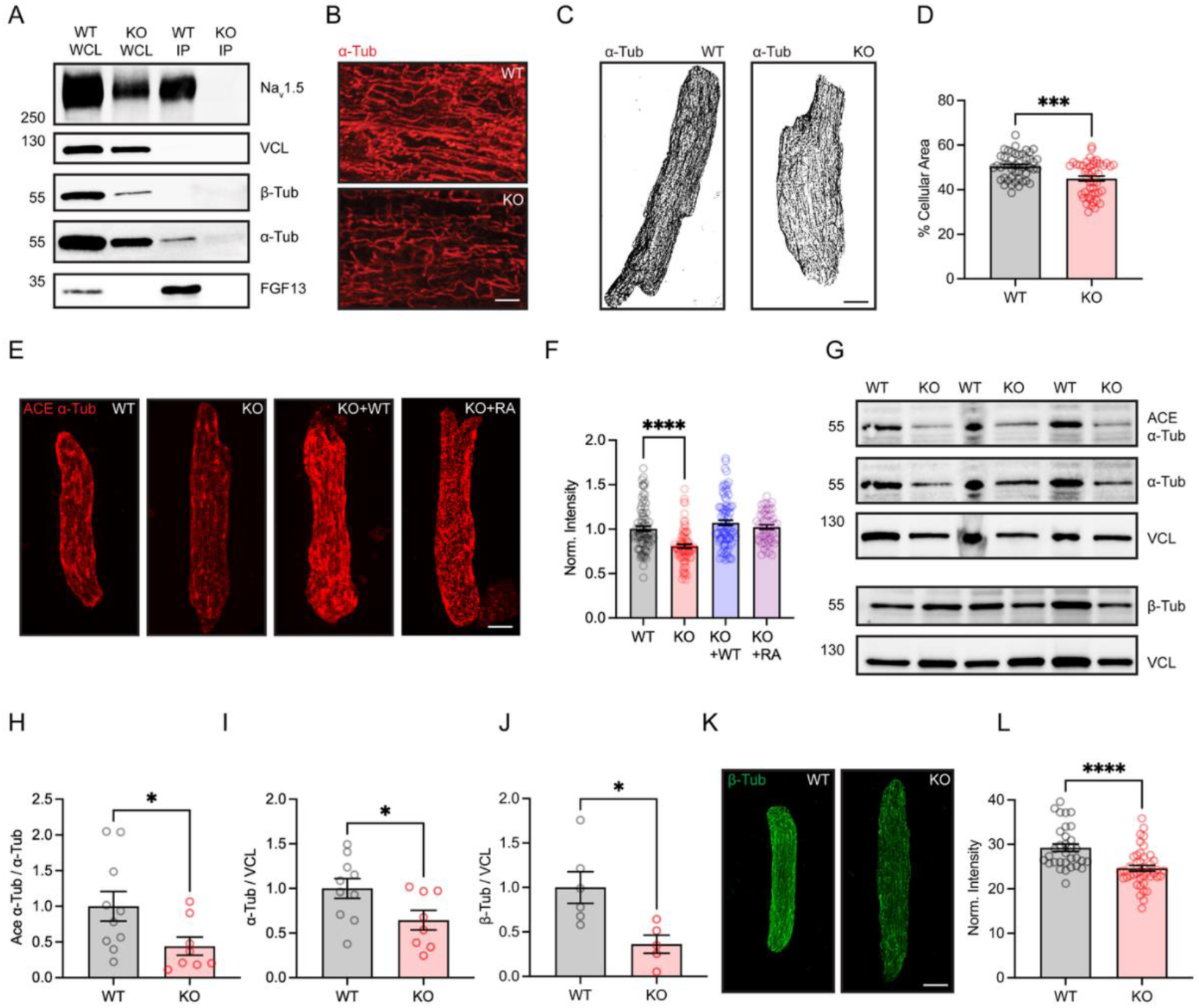
Loss of FGF13 reduces microtubule stability and tubulin expression. **A.** Coimmunoprecipitation (Co-IP) of FGF13 shows α-tubulin (but not β-tubulin) in complex. Na_v_1.5 is a positive control. Vinculin (VCL) is a loading control. Protein concentration of IP fraction was 1/8^th^ of whole cell lysate (WCL). **B.** Representative 3D renderings of α-tubulin immunostain in WT and KO ventricular tissue sections. Scale bar, 10 µm. **C.** Representative images of tubulin-stained WT and KO myocytes, converted to 8-bit and thresholded. Scale bar, 15 µm. **D.** Summarized data of % Cellular Area of tubulin (*n =* 4 mice per group, WT *n =* 47 cells, KO *n* = 47 cells), *p* = 0.0002, 2-tailed, unpaired *t*-test with Welch’s correction. **E.** Representative images of WT, KO, KO + WT FGF13, and KO + FGF13^R/A^ re-expressed myocytes immunostained with acetylated α-tubulin. Scale bar, 10 µm. **F.** Summarized data of acetylated α-tubulin (ACE) mean fluorescence (pixel) intensity (*N* = 4 mice per group, WT *n* = 80 cells, KO *n* = 78 cells, KO + WT FGF13 *n* = 83 cells, KO + FGF13^R/A^ *n* = 51 cells), one-way ANOVA corrected for multiple comparisons, *p* < 0.0001 for WT vs KO. **G.** Representative immunoblots of ACE, α-tubulin, β-tubulin, and VCL (*N* = 3 mice per group). **H**. Summarized relative expression densitometric data of ACE normalized to α-tubulin from immunoblots (WT *N =* 10 mice, KO *N* = 8 mice), 2-tailed, unpaired *t*-test with Welch’s correction, *p* = 0.036. **I.** Summarized relative expression densitometric quantification of α-tubulin normalized to VCL (WT *N =* 10 mice, KO *N* = 8 mice), 2-tailed, unpaired *t*-test with Welch’s correction, *p* = 0.036. **J.** Representative immunoblots β-tubulin and VCL (*N* = 3 mice per group). **K.** Summarized relative expression densitometric quantification of β-tubulin normalized to VCL (WT *N =* 6 mice, KO *N* = 5 mice), 2-tailed, unpaired *t*-test with Welch’s correction, *p* = 0.015. **L.** Representative confocal images of acutely isolated WT and KO ventricular myocytes immunostained with β-tubulin. Scale bar, 15 µm. **M.** Summarized data of β-tubulin mean fluorescence (pixel) intensity (*N* = 3 mice per group, WT *n* = 32 cells, KO *n* = 42 cells), 2-tailed, unpaired *t*-test with Welch’s correction, *p* < 0.0001. \**p* < 0.05, \*\*\**p* < 0.001, \*\*\*\**p* < 0.0001.

The overall regulation of tubulins and microtubules by FGF13 in cardiomyocytes prompted us to investigate the role of MAP4, a critical modulator of microtubule stability in cardiomyocytes, a key microtubule-associated protein implicated in cardiac health and disease ^37,38^ and a protein identified in the TurboID screen as a candidate FGF13 near neighbor (**Supplemental Figure 7C**). We therefore assessed MAP4 in WT and c*Fgf13^KO^* hearts (**Figure 8A-B**) and in isolated cardiomyocytes (**Figure 8C-D**) and found that MAP4 abundance was significantly reduced in the knockout hearts. Additionally, *Map4* mRNA was reduced in c*Fgf13^KO^* hearts (**Supplemental Figure 5C**). Overall, these results suggest that FGF13-deficiency in ventricular myocytes reduces the abundance of MAP4 and tubulin as well as markers of microtubule stability, and perturbs trafficking of Cx43.

**Figure 8.**
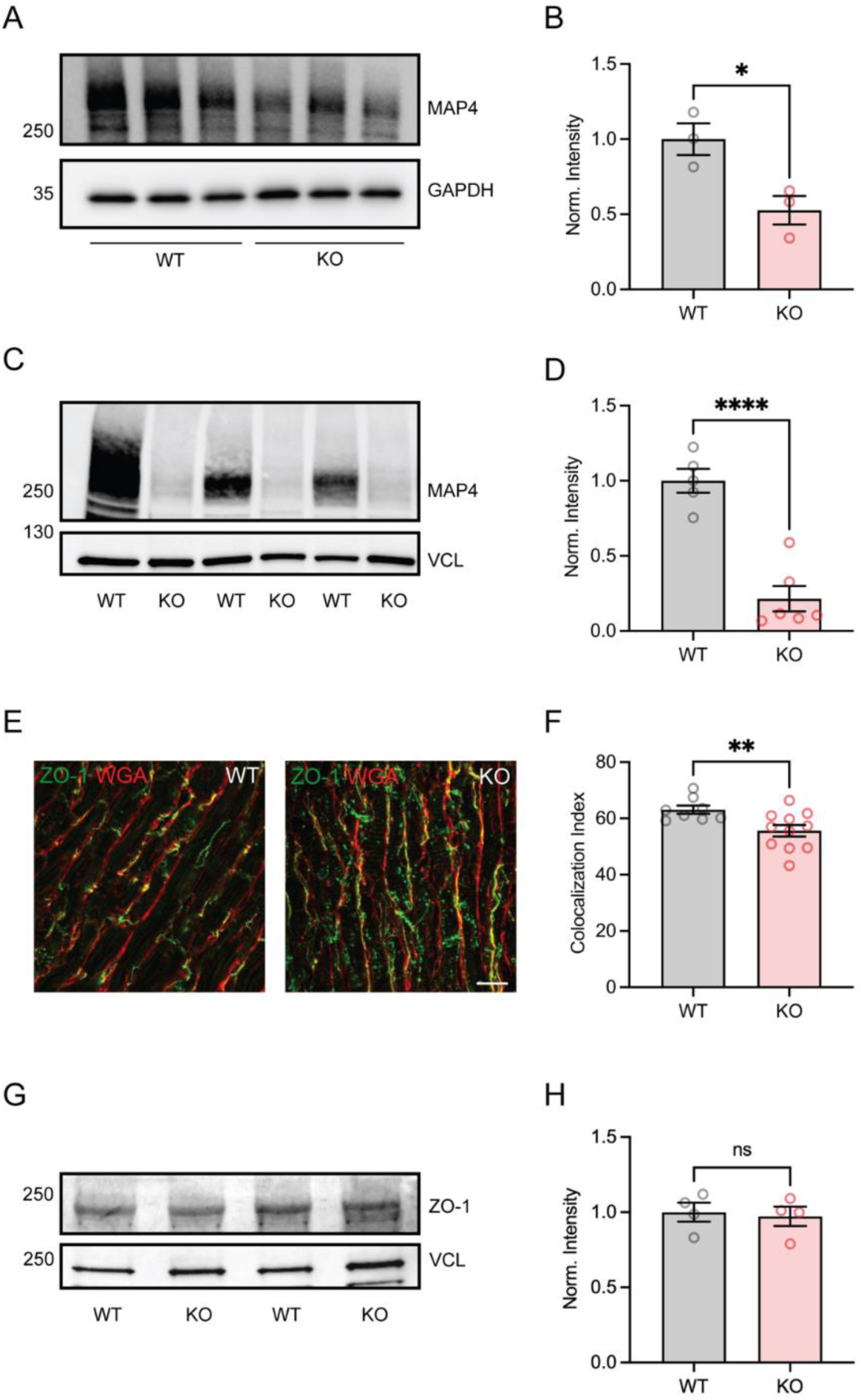
Constitutive loss of FGF13 decreases MAP4 expression as well as impacts trafficking of ZO-1. **A.** Western blot of MAP4 and GAPDH from WT and c*Fgf13^KO^* whole heart lysates (*N* = 3 mice per group). **B.** Densitometric analysis of whole heart MAP4 protein expression (from **A**) normalized to GAPDH (*N* = 3 mice per group), 2-tailed, unpaired *t*-test with Welch’s correction, *p* = 0.0293. **C.** Western blot of MAP4 and Vinculin from WT and c*Fgf13^KO^* isolated ventricular myocyte lysates (*N* = 3 mice per group). **D.** Densitometric analysis of MAP4 protein expression from isolated myocyte lysates normalized to vinculin (WT, *N* = 5 mice; KO, *N* = 6 mice), 2-tailed, unpaired *t*-test with Welch’s correction. **E.** Representative images of WT and c*Fgf13^KO^* ventricular cryosections immunostained for ZO-1 (green) and WGA (red). Scale bar, 20 µm. **F.** Quantification of colocalization between ZO-1 and WGA in tissue sections (*N =* 3 mice per group, 2-3 sections per mouse (WT *n* = 8 fields, KO *n* = 10 fields), *p* = 0.008, 2-tailed, unpaired *t-* test with Welch’s correction. **G.** Western blot of ZO-1 (Vinculin as a loading control). **H.** Densitometric analysis of ZO-1 protein expression from isolated myocyte lysates normalized to vinculin (*N* = 4 mice, each). \**p* < 0.05, \*\**p* < 0.01, \*\*\**p* < 0.001, \*\*\*\**p* < 0.0001.

Since multiple proteins rely on vesicular transport via microtubules to reach their specific subcellular destinations, we hypothesized that FGF13 knockout affected the trafficking and targeting of proteins other than Cx43. In the FGF13 TurboID screen (**Figure 4** and **Supplemental Dataset 1**), we identified multiple sarcolemmal ion channels and other ID proteins (**Supplemental Table 1**). Among these was the tight junction protein ZO-1 (*Tjp1*, Log_2_FC=4.20, −log_10_P_adj_=2.95), a component of the ID and regulator of Cx43 organization ^39^. We performed immunocytochemistry for ZO-1 and found that, similar to Cx43, ZO-1 distribution was perturbed (**Figure 8E-F**). However, unlike Cx43, ZO-1 expression was similar between c*Fgf13^KO^*and WT hearts (**Figure 8G-H**).

## DISCUSSION

Although FGF13 is best known and characterized as a regulator of VGSCs through FGF13’s direct interaction with their C-termini ^5,7^, several reports suggest that FGF13 has additional roles beyond voltage-gated channel regulation. Among these, we previously determined that FGF13 regulates the abundance of K^+^ channels at the sarcolemma ^16^. With results here showing that FGF13 also directs the trafficking and targeting of Cx43 and ZO-1, we suggest that FGF13 is a regulator of trafficking and targeting of multiple proteins utilizing microtubule-dependent vesicular transport. Intriguingly, we found no difference in *Cx43* transcription or Cx43 degradation despite the overall increased Cx43 abundance in c*Fgf13^KO^*cardiomyocytes. These results suggest that FGF13 may affect the rate or efficiency of *Cx43* translation. Indeed, gene ontology for the TurboID dataset of terms found in the FGF13 near neighborhood but not in the Na_V_1.5 near neighborhood (**Figure 6E**) identifies “regulation of translation” among the top biological process terms (GO:0006417, *P*_adj_ = 3.77E-15).

While previous studies postulated that FGF13 affects Na_V_1.5 trafficking and targeting via FGF13’s direct interaction with the Na_V_1.5 C-terminus, results here suggest that FGF13’s trafficking and targeting functions are more general. Indeed, we recently exploited the mutant FGF13^R/A^, which is unable to bind to Na_V_1.5, and showed that FGF13^R/A^ restored critical aspects of VGSC localization in c*Fgf13^KO^* cardiomyocytes via regulation of local membrane cholesterol ^22^. Thus, even some components of VGSC regulation by FGF13 are independent of FGF13 binding to the VGSC C-termini. The ability of FGF13^R/A^ to restore trafficking and targeting of Cx43 in this study suggests that FGF13’s effects on Cx43 are also independent of regulation of VGSC by FGF13.

The mechanisms by which FGF13 affects protein trafficking and targeting and thus increase the susceptibility of c*Fgf13^KO^* mice to arrhythmias in the setting of Cx43 blockade rely on microtubules and may include several independent regulatory steps. Specifically, we showed that FGF13 is in complex with, and regulates, microtubules in cardiomyocytes as well as controls the abundance of tubulins and the key cardiomyocyte microtubule regulator MAP4. Whether the reduction in MAP4 is upstream of microtubule instability or the reduction in tubulins and acetylated tubulins drives the decrease in MAP4 is not known. We note that the microtubule deacetylases (HDAC6, SIRT2) were enriched in the FGF13 TurboID screen, and thus FGF13-dependent regulation of their activity may also contribute to the observed decrease in alpha tubulin acetylation in c*Fgf13^KO^*hearts and consequent microtubule stability. Our results suggest that the microtubule instability in the absence of FGF13, affects trafficking and targeting of Cx43 and other components of the ID. This leads to reduced Cx43 at the ID and increased Cx43 hemichannels at the lateral membrane, thus providing the substrate for arrhythmogenesis in the c*Fgf13^KO^* mice. Our results showing that Cx43 subcellular localization is perturbed in c*Fgf13^KO^* cardiomyocytes and that increased Cx43 hemichannels contribute to slowed conduction velocity appear to contrast with a previous study that reported no difference in the localization of Cx43 in *Fgf13* knockout myocytes ^12^. That whole-body knockout model did report that *Fgf13* knockout hearts were more sensitive to the gap junction inhibitor carbenoxolone, consistent with reduced Cx43 reserve. The reasons for the study differences are not clear, but we suspect that they reflect different levels of *Fgf13* reduction. Others ^17^ and we ^18,19^ observed that complete *Fgf13* knockout animals are nonviable ^15,17^, in contrast to the *Fgf13* whole body knockout study ^12^. That model may thus be a hypomorph, leaving residual FGF13 expression. Consistent with the modeling data from that study, we also conclude that the failed impulse propagation with *Fgf13* deficiency does not result only from a reduction in VGSC conductance. Results in **Figure 1H** show slowed CV at all pacing rates tested, even at slower rates at which VGSC inactivation is minimized, thus implicating mechanisms beyond reduced VGSC conductance ^40^. At least one contributor appears to be the depolarized RMP in c*Fgf13^KO^* cardiomyocytes that would drive more VGSC inactivation due to the relatively steep steady-state inactivation curve near RMP ^22^. Further, our results suggest that underlying the depolarized RMP in the context of *Fgf13* deficiency is an increase in Cx43 hemichannels as shown by the restoration of RMP to WT levels with the Cx43 hemichannel blocker Gap19 (**Figure 4G**). Cx43 hemichannels were recently recognized as contributors to inward depolarizing currents in disease states such as heart failure ^23,25^, suggesting investigations into whether perturbation of FGF12, the homologous FHF predominant in human heart, contributes to arrhythmogenesis in heart failure.

In summary, our results suggest broader effects of FGF13 than regulation of VGSCs in cardiomyocytes and imply that multiple channels and auxiliary subunits important for normal rhythm are perturbed in *Fgf13* knockout cardiomyocytes. These results add to recent observations about the non-VGSC role for FGF13 in regulation of neuronal excitability ^19^, and emphasize the importance of exploring additional mechanisms to understand how FHFs contribute to cardiac physiology and how variants in FHFs lead to pathophysiology.

## Methods

### FGF13 constitutive knockout animals

To generate cardiac-specific constitutive knockout mice (c*Fgf13^KO^*) of the X-linked *Fgf13*, we crossed female *Fgf13^fl/fl^* ^16^ with hemizygous male *Myh6-Cre* (alpha-Myh6-Cre; The Jackson Laboratory, #011038) mice to generate hemizygous male knockout animals. Mice were maintained on the C57BL/6 background. Experiments were performed on 6-to 12-week-old mice.

### Generation of cardiac-specific TurboID-FGF13VY transgenic mice

The TurboID-hFGF13VY construct was generated by fusing the cDNA of TurboID ^32^, containing a Hemagglutinin (HA) tag and a V5 tag at the N-terminal domain of the hFGF13VY cDNA. The construct was then cloned into the modified murine α-myosin heavy chain (MHC) tetracycline-inducible promoter vector ^41,42^ to generate TurboID-hFGF13VY transgenic mice with non-targeted insertions of the tetracycline-regulated transgene in a C57BL/6 background. These mice were crossed with cardiac-specific (α-MHC), doxycycline-regulated, reverse transcriptional trans-activator (rtTA) mice to generate double transgenic mice ^43^.

### Generation of cardiac-specific TurboID-SCN5A transgenic mice

The V5-TurboID-NaV1.5 knock-in model was generated by Cyagen. The Kozak-3X HA tag-TurboID-V5 tag (V5-TurboID) was created by gene synthesis and inserted upstream of the ATG start codon, which is in exon 2 of the *Scn5a* gene located on mouse chromosome 9. The targeting vector included homology arms generated by PCR using BAC clone RP23-386N9 or RP23-198L19 with V5-TurboID and a Neo cassette flanked by self-deletion anchor (SDA) sites. Following successful targeting, diphtheria toxin A was employed for negative selection. The targeting construct was electroporated into embryonic stem cells. Positive clones were identified by PCR and confirmed by Southern blotting. Five clones were injected in C57BL/6N blastocysts, and chimeras were crossed to C57BL/6 mice. Heterozygous mice were bred to produce homozygous offspring.

### *In vivo* biotinylation

Male and female double transgenic mice, 4-8 months old, were placed on a doxycycline diet (in pelleted form, 200mg/kg of diet) for 10 days to express the TurboID-FGF13VY transgene in the heart. Biotin (10 µl/g from a 2.4 mg/ml stock (in PBS:DMSO 9:1) or 10% DMSO in PBS (for the controls)) were injected daily, subcutaneously, for 3 consecutive days to achieve biotinylation in the hearts of TurboID-FGF13VY and V5-TurboID-Na_V_1.5. The mice were sacrificed to collect the hearts 24 hours after the third injection. Whole heart tissues were lysed with a handheld tip homogenizer in Lysis Buffer (50 mM Tris-HCl pH 7.5, 150mM NaCl, 10mM EDTA, 0.5% Sodium Deoxycholate (m/v), 1% Triton X-100 (v/v), 0.1% SDS (w/v)) supplemented with Roche cOmplete Protease Inhibitor Cocktail and 1 mM PMSF (PhenylMethylSulfonyl Fluoride). Biotinylation efficiency was confirmed by Western blot analysis with Streptavidin-HRP.

### Sample preparation for mass spectrometry

Heart tissue samples for mass spectrometry analysis were prepared as previously described ^44^ and adapted for this study. Whole heart lysates were centrifuged at 21,130 x g at 4 °C for 15 minutes. 1 mg was precipitated with ice-cold trichloroacetic acid (TCA, 55% in double-distilled water) on ice for 15 minutes and centrifuged at 21,130 x g at 4 °C for 15 minutes. The pellets were washed with −20 °C cold acetone, resuspended and centrifuged at 21,130 × g at 4 °C for 10 minutes. Acetone was removed and the pellets washed with acetone again three times to ensure complete TCA removal. After the last acetone wash the pellets are resuspended in urea buffer (8M urea, 100 mM sodium phosphate pH 8, 100 mM NH_4_HCO_3_, and 1% SDS (w/v), centrifuged at 21,130 x g at room temperature for 10 minutes, transferred to new microcentrifuge tubes and 10mM Tris(2-carboxyethyl)phosphine was added to reduce disulfide bonds. Twenty mM of freshly prepared iodoacetamide was added to alkylate free cysteines, vortexed immediately and incubated for 25 minutes in the dark. Alkylation was then quenched by adding 50mM of dithiothreitol (DTT) and water was added to dilute the samples down to 4M urea and 0.5% weight/volume (w/v) sodium dodecyl sulfate (SDS). Samples were added to 50 µl of streptavidin magnetic beads pre-washed in 4 M urea, 100 mM sodium phosphate pH 8, 0.5% w/v SDS and tubes were rotated overnight at 4 °C to for streptavidin pull-down. The beads were then transferred to a new microcentrifuge tube, washed 3 times with 4 M urea, 100 mM sodium phosphate pH 8, 0.5% SDS (w/v) three times with 4 M urea, 100 mM sodium phosphate pH 8 without SDS. The beads were transferred to new tubes for one last wash step. Before the final bead pull-down 10% of the resuspended beads were collected for streptavidin-HRP blotting.

### Mass spectrometry analysis

Mass spectrometric analysis for the FGF13 TurboID dataset was performed at the Weill Cornell Medicine Proteomics and Metabolomics Core Facility. In brief, the protein on beads were reduced with DTT, alkylated with iodoacetamide, and digested overnight with trypsin at 37 °C. The digests were desalted by C18 Stage-tip columns. The digests were analyzed using a Thermo Fisher Scientific EASY-nLC 1200 coupled on-line to a Fusion Lumos mass spectrometer (Thermo Fisher Scientific). Buffer A (0.1% FA in water) and buffer B (0.1% FA in 80 % ACN) were used as mobile phases for gradient separation. A 75 µm x 15 cm chromatography column (ReproSil-Pur C18-AQ, 3 µm, Dr. Maisch GmbH, German) was packed in-house for peptide separation. Peptides were separated with a gradient of 5–40% buffer B over 30 min, 40%-100% B over 10 min at a flow rate of 400 nl/min. The Fusion Lumos mass spectrometer was operated in a data independent acquisition (DIA) mode. MS1 scans were collected in the Orbitrap mass analyzer from 350-1400 m/z at 120K resolutions. The instrument was set to select precursors in 45 x 14 m/z wide windows with 1 m/z overlap from 350-975 m/z for HCD fragmentation. The MS/MS scans were collected in the Orbitrap at 15K resolution. Data were searched against the mouse Uniprot database (downloaded on August 7, 2021) using DIA-NN v1.8 and filtered for 1% false discovery rate for both protein and peptide identifications. Statistical significance was determined by multiple t-tests. Proteomic analysis for data of FGF13-TurboID is displayed in **Figure 6** and **Supplemental Dataset 1.** TMT quantified proteomic analysis for data of Na_V_1.5-TurboID as displayed in **Figure 6** and in **Supplemental Dataset 2** was performed as previously described for APEX-fusion proteins following an SPS-MS3 mass spectrometry method ^45^.

For Gene Ontology (GO) term analysis, proteins with a Log2(biotinylated/ctrl)>2, P_adj_ < 0.05 were used and g:Profiler ^46^ was used to generate GO term tables. For each GO term, we first computed the Gene Ratio by dividing the Intersection_Size value with the Query_Size value. We then divided the Gene Ratio by (Term_Size / Effective_domain_size) to derive the Enrichment Ratio. We sorted the GO terms by the product of -log_10_P_adj_ and the Enrichment Ratio and displayed the top 13 terms.

### Electrocardiograms

Mice were anesthetized with tribromoethanol (Avertin) and single lead ECGs were recorded subcutaneously, while their body temperature was maintained with a heat lamp at 37 °C. Signals were amplified and recorded with an Octal Bio Amp amplifier connected to a Powerlab 16/30 DAQ system (ADInstructions). Heart rate, PR, QRS and QT intervals were analyzed using LabChart, version 8.0, software (ADInstruments). The ECG Analysis Module for LabChart, with the following specifications, was used for interval analysis: Typical QRS width 10 ms, R waves > 60 ms apart, and Pre-P baseline 10 ms. Fifty to 100 beats were selected and analyzed with 4 beat averaging. Automated outputs were manually checked for accuracy using the ECG Averaging View and edited as needed. Conduction block was determined when there was no QRS complex after a P wave. For ECG recordings with carbenoxolone, carbenoxolone disodium salt (C4790, Sigma-Aldrich) was dissolved in distilled water (90 mg/kg) and injected intraperitoneally. Surface ECGs were recorded 30 minutes after carbenoxolone administration for 15 min.

### Electrophysiology Recordings

Action potentials were recorded in an extracellular solution containing (in mM): 140 NaCl, 5.4 KCl, 1.8 CaCl₂, 1 MgCl₂, 10 glucose, and 10 HEPES, with pH adjusted to 7.4 with NaOH. Patch pipettes made from borosilicate glass (2–2.5 MΩ resistance) were filled with an internal solution consisting of (in mM): 125 K-gluconate, 20 KCl, 5 NaCl, 1 MgCl₂, 5 MgATP, and 10 HEPES, adjusted to pH 7.2 with KOH. All chemicals were purchased from Sigma-Aldrich. To compare action potentials between WT and KO myocytes, the perforated patch technique was employed with 200 μg/ml amphotericin B added to the internal solution and action potentials were elicited by injecting a 200 pA current pulse for 20 ms. For all other action potential recordings, the whole-cell patch clamp technique was used. In the FGF13 rescue experiments, cells were held at a membrane potential of −90 mV, and action potentials were recorded at rheobase. In GAP19 treatment experiments, cells were held at their resting membrane potential, and action potentials were induced by injecting a 200 pA current pulse for 50 ms.

### Optical Mapping

Mice (8-12 wk old) were injected with 0.3 ml of a 1:1 heparin:saline solution and euthanized. Hearts were rapidly excised, cannulated, and perfused with oxygenated Tyrode’s solution (in mM: 129 NaCl, 24 NaHCO_3_, 4 KCl, 1 MgCl_2_, 11.2 mM glucose, 1.8 CaCl_2_, 1.2 KH_2_PO_4_) on a Langendorff apparatus. The perfusate contained Blebbistatin (10 µM, Sigma-Aldrich, #B0560) to suppress motion artifacts and 30 µl of 2.5 mM Di-4-ANEPPS (ThermoFisher #D1199) was injected via inline port. Hearts were submerged in a 37 ± 1°C bath of Tyrode’s with perfusion pressure maintained at ∼70 ± 15 mm Hg by adjusting flow (2.0 ± 1 ml/min).

A pacing electrode was placed in the left ventricular apex. ECG and high-resolution optical signals of the anterior epicardial surface of the ventricles were acquired as previously described with a 6×6 mm² field of view ^47^. Diastolic threshold was determined at a pacing rate of 7.14 Hz; experiments were run at 1.5× diastolic threshold.

The hearts were subjected to a rapid pacing protocol under normal conditions and then the perfusate switched to low potassium Tyrode’s (2 mM KCl). The hearts were perfused with low potassium Tyrode’s for 20 min before rapid pacing protocol was repeated.

#### Rapid Pacing Protocol

Pacing rate progressively increased from 8 to 24Hz with two-second recordings taken at each step. If there was loss of capture, voltage was increased until capture regained, or maximum voltage was hit with no capture and the protocol was ended. After switching to new perfusate, the voltage was reset to 1.5x diastolic threshold before beginning the rapid pacing.

#### Optical Mapping Analysis

Optical recordings were analyzed using the open-source software ElectroMap in MATLAB ^48^. The ventricles were manually traced as the region of interest. Spatial and signal processing settings included a 3×3 pixel Gaussian filter (sigma=1.5), Savitzky-Goaly filter, and Top-Hat average filter (length of 100) for baseline correction. For each pacing rate, 10 consecutive beats were analyzed, and conduction velocity (CV) was calculated through ensemble averaging. Data were graphed in GraphPad Prism 10.

### Cardiomyocyte Isolation, Cell Culture and Adenoviral Transduction

Adult murine cardiomyocytes were isolated from left and right ventricles of Myh6-Cre^+^ (WT) and *Myh6-Cre+;Fgf13^fl/fl^* (cFGF13KO) male mice using a previously validated Langendorff-free cell isolation method ^49^. Briefly, mice aged 8-12 weeks were anesthetized with Avertin and the chest was incised to expose the heart. The inferior vena cava and descending aorta were cut, and the heart was flushed by injecting 10 ml of room temperature EDTA buffer into the right ventricle. EDTA buffer contains (in mM): 130 NaCl, 5 KCl, 0.5 NaH_2_PO4, 10 HEPES, 10 Glucose, 10 2,3-butanedione 2-monoxime (BDM), 10 Taurine, 5 EDTA. After placing an aortic clamp, the heart was excised and transferred to a 60-mm dish containing EDTA buffer. Next, 10 ml of room temperature EDTA buffer was injected into the apex region of the left ventricle. The same aperture was used to inject 5 ml of perfusion buffer (warmed to 37C) and then serial injection of collagenase buffer (5X 10 ml tubes with collagenase buffer) into the left ventricle (LV). Perfusion buffer contains (in mM): 130 NaCl, 5 KCl, 0.5 NaH_2_PO_4_, 10 HEPES, 10 Glucose, 10 BDM, 10 Taurine, 1 MgCl_2._ The atria were then separated and discarded. The ventricles were pulled into 1-mm pieces with forceps and cells were dissociated with gentle trituration for 2 minutes. Enzymatic activity was inhibited by addition of 3 ml of stop buffer (10% FBS in Perfusion Buffer), and the cell suspension was passed through a 150-mm filter and allowed to gravity settle for 20 minutes. The cells were then washed twice in perfusion buffer and allowed to gravity settle for 10 minutes each and used for various experiments.

Cell culture and adenoviral infection of adult mouse ventricular myocytes were performed as described ^2,49^. Briefly, prior to cardiomyocyte isolation, 12-well plates were coated with laminin at a concentration of 5 mg/ml in PBS, for 1 hour at 37°C. After isolated cardiomyocytes were retrieved from gravity settling, the laminin solution was aspirated, the wells were washed with PBS, and cardiomyocytes equilibrated in Plating Media (described in ref. ^49^) were added to the wells. Cells were allowed to adhere for 1 hour. Subsequently, the Plating Media was removed, and the cells were incubated in 500 ml of Culture media per well. After 2 hours in Culture media described in ref. ^49^), a subset of KO cardiomyocyte wells was infected with 1.5 µl of FGF13-VY adeno-associated virus serotype 8 (AAV8, 1.6 X 10^8^ ifu/ml) or 1.5 µl of FGF13^R/A^ AAV8 (6.6 X 10^7^ ifu/ml). The control group was infected with an AAV8 expressing GFP. Cardiomyocytes were incubated in Culture Media for 48 hours for adequate viral expression before subsequent analyses were performed. Electrophysiological analyses were obtained from recording sessions in which all relevant experimental groups had been simultaneously cultured. The amount of AAV-expressed FGF13 or FGF13^R/A^ in the c*Fgf13^KO^* cells was similar to the level of endogenous FGF13, as shown previously ^22^.

### Immunoblots

Isolated cardiac myocytes were lysed for 30 minutes in ice cold RIPA buffer supplemented with Halt protease and phosphatase inhibitor cocktails (Thermo Fisher Scientific). The homogenate was then centrifuged at 17,000 g at 4C for 15 minutes. Supernatant was collected and protein concentration was determined by bicinchoninic acid assay. Protein was separated on 8-16% gradient Tris-Glycine gels (Thermo Fisher Scientific) and transferred to PVDF membranes using iBlot 3 Western Blot Transfer System (Thermo Fisher Scientific) in Broad Range mode for 6 minutes. Membranes were immunoblotted for Na_V_1.5 (1:1000, 493-511, Alomone Labs), FGF13 (1:1000, custom designed ^2^) GAPDH (1:1000, MA5-15738, Thermo Fisher Scientific), Cx43 (1:1000, C6219, Sigma-Aldrich), ZO-1 (1:500, 5406, Cell Signaling Technology), vinculin (1:1000, sc-73614, Santa Cruz Biotechnology), Na^+^/K^+^ ATPase (1:1000, sc-48345, Santa Cruz Biotechnology), α-tubulin (1:1000, sc-5286, Santa Cruz Biotechnology), β-tubulin (1:1000, 2146, Cell Signaling Technology), acetylated α-tubulin (1:1000, 5335S, Cell Signaling Technology), and MAP4 (1:500, 11229-1-AP, Proteintech). All primary antibodies were incubated overnight at 4 °C at respective concentrations. HRP-tagged secondary antibodies were incubated at room temperature for 1 hour. The blots were visualized by chemiluminescence, and images were captured using ChemiDoc Tough Imaging System (Bio-Rad).

### Co-immunoprecipitation

Co-immunoprecipitation shown in Figure 2A was performed as following: hearts from 3 WT mice, 4-6 months old, were collected and lysed in NP40 Lysis buffer (150mM NaCl, 20mM Tris-HCl pH 8, 2mM EDTA, 1% NP40 (v/v)) supplemented with Roche cOmplete Protease Inhibitor Cocktail and 0.2 mM PMSF. One mg of lysate was pre-cleared with Dynabeads Protein A beads (Thermo Fisher Scientific) for 1 h at 4 °C. Pre-cleared lysates were incubated with 5 µl of Nav1.5 antibody (Alomone #493-511) (0.8 µg/µl, Alomone Labs) overnight at 4 °C. Immunoprecipitation was performed with 50 µl of Dynabeads Protein A beads for 1 h at room temperature and proteins were eluted with 50 µl of 0.2 M glycine at pH 2.4 for 15 min at room temperature, immediately neutralized with an equal volume of 1 M Tris-HCl, pH 8.

Co-immunoprecipitation for Figure 7 was performed according to the manufacturer’s instruction (Capturem IP & Co-IP Kit, Takara, 635721). Briefly, isolated ventricular myocytes (nearly 5X 10^5^ per genotype) were lysed on ice for 30 min with 200 µl lysis buffer. Following 17,000 X g centrifugation at 4C for 10 min, the supernatant was incubated with 10 µg antibody (maintaining an antibody: lysate concentration of 2:500) for 20 minutes at room temperature. After incubation, the antibody-bound lysate was added onto the spin column and centrifuged at 1000 X g for 1 min at room temperature. Then, 100 µl wash buffer was added to the spin column and centrifuged at 1000 X g for 1 min again. Finally, 50 µl elution buffer was added to the spin column and centrifuged at 1000 X g for 1 min. The eluted sample was used for Western blot analysis as previously described ^33^.

### Quantitative Real-Time PCR

RNA was extracted from isolated ventricular cardiomyocytes using instructions from the RNeasy Mini Kit (QIAGEN). Reverse transcription was performed using Bio-Rad’s instructions from the iScript cDNA Synthesis Kit. For each sample, qPCR was performed in duplicate using SYBR green-based detection (Bio-Rad). Ct values were quantified using Quantstudio 3 (Applied Biosystems). *Gapdh* or *Vcl* were used as a reference gene. PCR primer sequences for target genes are as follows:

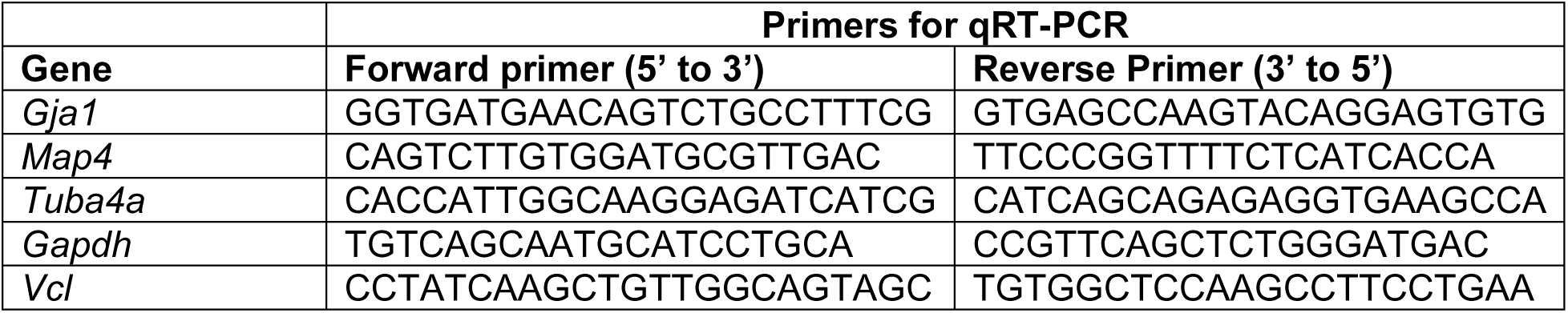

### Histological Analysis

Hearts were rinsed in PBS, fixed in 4% paraformaldehyde overnight at 4C, and dehydrated in a series of ethanol washes. Samples were cleared in xylene and mounted in paraffin. Sections of 10 µm were cut and stained with hematoxylin and eosin to assess gross morphology. Samples from different groups were processed in parallel, and histological analyses performed. Immunostaining and confocal microscopy were performed as described ^50^.

### Echocardiography

Echocardiography was performed on conscious mice by a technician blinded to animal genotype as previously described ^15^. In brief, images were obtained on a VEVO 2000 high-resolution imaging system (VisualSonics). Transthoracic 2D M-mode used for data analysis. Heart rate, left-ventricular end-diastolic diameter (LVEDD), and left-ventricular end-systolic diameter (LVESD) were measured from at least three consecutive cardiac cycles by two experimenters blinded to genotype. Fractional shortening (FS) was calculated with the formula: FS = (LVEDD – LVESD)/LVEDD.

### Quantitative Immunocytochemistry Analysis

Isolated cardiomyocytes were fixed for 10 minutes in 4% paraformaldehyde and immunolabeled with antibodies against Cx43 (C6219, Sigma, St Louis, MO) and N-cadherin (sc-59987, Santa Cruz Biotechnology). Images were captured on a Zeiss LSM 800 confocal microscope and the amount of immunoreactive signal at the ICD was analyzed with NIH ImageJ/FIJI software as described previously ^51^. In addition, using a Pearson-Spearman correlation colocalization plug-in, quantitative statistical colocalization on the two-color confocal images was performed as described ^52^. For ZO-1-WGA colocalization image analysis, hearts from 2-month-old WT and FGF13 KO mice were harvested, rinsed in PBS, and incubated sequentially in 15% and 30% sucrose solutions. Hearts were then incubated in OCT medium (TissueTek) for 1 hour and then frozen in OCT. Ten micron cryosections were made, permeabilized with 0.1% Tween-20 + PBS (PBST), and blocked in PBST + 2% Fetal Bovine Serum (FPBST) for 1 hour. Primary antibodies were incubated overnight with respective secondary Alexa-Fluor antibodies for 1 hour at room temperature. After mounting, images were obtained at 20X using Thunder Leica Microscope. In ImageJ, the red (WGA) and green (ZO-1) channels were converted to 8-bit images. The ZO-1 mean pixel intensity per image was measured. Then the WGA intensity was subtracted from ZO-1 and the residual pixel intensity was measured. The residual intensity indicated the amount of non-colocalized ZO-1. After initial optimization and development of analytical algorithms, the experimenter was blinded to genotype for image analyses.

### Biotinylation

Biotinylation was performed according to the manufacturer’s instruction (Pierce^TM^ Cell Surface Biotinylation and Isolation Kit, ThermoFisher Scientific, A44390).

### Area and Pixel Intensity Calculations

Isolated cardiomyocytes were fixed in 4% paraformaldehyde for 10 minutes and Cx43 was identified with anti-Cx43 antibody (C6219, Sigma, St Louis, MO). Images were captured in 40X using a Leica immunofluorescence microscope. The images were opened in NIH/ImageJ software, converted to 8-bit images, and a black and white threshold was applied. Then the cell border was manually drawn, and the area was measured as a percentage of the cell. For pixel intensity calculations of confocal images, a max intensity z-stack was generated. Then the background was subtracted. A line was drawn along the longest axis of the cell, and pixel distribution generated as a Plot Profile. For calculation of Cx43 puncta size from confocal images, a max intensity z-stack was generated. In ImageJ/FIJI, each image’s background was subtracted, converted to 8-bit, and thresholded. The lower threshold was set to 22 (based on clear visibility of small and large puncta with minimal pixel overlap) and applied to all images. The number of particles was counted using the “Analyze Particles” function. Small puncta were defined as between 0 - 0.2 µm^2^ (based on literature showing that Cx43 loaded vesicles are <150 nm in diameter) and large puncta were counted between 0.21 - 1.80 µm^2^.

### Statistical Analysis

Results are presented as mean ± standard error; the statistical significance of differences between groups was assessed using either a two-tailed Student *t* test or one-way analysis of variance and was set at *P*<0.05. Other statistical analyses were performed as previously described ^15^ or as described in corresponding figure legends. No animals were excluded from analyses.

### Study approval

This study was approved by the Weill Cornell Institutional Animal Care and Use Committee (Protocol #. 2016-0042) and the Yale Institutional Animal Care and Use Committee (Protocol # 2023-20360. All animals were handled in accordance with the NIH *Guide for the Care and Use of Laboratory Animals*. All mice were maintained on a C57BL/6 genetic background.

